# ABHD4-dependent developmental anoikis protects the prenatal brain from pathological insults

**DOI:** 10.1101/2019.12.17.879551

**Authors:** Zsófia I. László, Zsolt Lele, Miklós Zöldi, Vivien Miczán, Fruzsina Mógor, Gabriel M. Simon, Ken Mackie, Imre Kacskovics, Benjamin F. Cravatt, István Katona

**Affiliations:** Momentum Laboratory of Molecular Neurobiology, Institute of Experimental Medicine, Budapest, Hungary; School of Ph.D. Studies, Semmelweis University, Budapest, Hungary; Faculty of Information Technology and Bionics, Pázmány Péter Catholic University, Budapest, Hungary; The Skaggs Institute for Chemical Biology, Department of Chemical Physiology, The Scripps Research Institute, La Jolla, California, United States; Gill Center for Biomolecular Science, Department of Psychological and Brain Sciences, Indiana University, Bloomington, Indiana, United States; Department of Immunology, Eötvös Loránd University, Budapest, Hungary; ImmunoGenes Ltd, Budakeszi, Hungary

## Abstract

In light of the astronomical number of cell divisions taking place in restricted neurogenic niches, brain malformations caused by ectopic proliferation of misplaced progenitor cells are surprisingly rare. Here, we show that a process we term developmental anoikis distinguishes the abnormal detachment of progenitor cells from the normal delamination of daughter neuroblasts in the developing mouse neocortex. By using *in vivo* gain-of-function, loss-of-function, and rescue manipulations together with correlated confocal and super-resolution imaging, we identify the endocannabinoid-metabolizing enzyme abhydrolase domain containing 4 (ABHD4) as an essential mediator for the elimination of abnormally detached cells. Consequently, rapid ABHD4 downregulation is necessary for delaminated daughter neuroblasts to escape from anoikis. Moreover, ABHD4 is required for fetal alcohol-induced apoptosis, but not for the well-established form of developmentally controlled programmed cell death. These results suggest that ABHD4-mediated developmental anoikis specifically protects the embryonic brain from the consequences of sporadic delamination errors and teratogenic insults.

---

In the developing brain, radial glia progenitor cells (RGPCs) spawn various cell types including intermediate progenitor cells, neurons, astrocytes, oligodendrocytes and ependymal cells^1–4^. Following asymmetric cell division at the ventricular surface, the self-renewed RGPCs remain anchored to their neighbors via cadherin-based adherens junctions^5, 6^. This adhesion complex not only serves as a structural stabilizer, but it is also involved in important signaling mechanisms regulating cell cycle, proliferation and differentiation, indicating that adherens junction assembly and disassembly are tightly coupled to cell fate decisions^7–9^. Accordingly, the non-RGPC-fated daughter cells need to break down their adherens junctions to delaminate from the ventricular wall and to migrate to their functional destinations along the radial scaffold of RGPCs^6, 9, 10^.

Appropriately timed delamination is critically important for the mature daughter neuroblasts to follow their normal migratory route^6^. In contrast, premature delamination and abnormal dispersion of RGPCs which retain their proliferative capacity at ectopic locations represents a serious risk for brain malformations, such as focal cortical dysplasia or periventricular heterotopia, potentially predisposing to various forms of intellectual disabilities and neurological deficits^11–14^. Thus, considering the enormous number of cell division events and subsequent delamination steps that are required for the generation of hundreds of billions of neurons and other cell types in the brain together with the remarkably low prevalence of congenital brain malformations, it is conceivable to hypothesize that a specific mechanism exists that protects against the consequences of sporadic delamination errors. Yet, how prematurely detached RGPCs are eliminated from the brain parenchyma and the related fundamental question of how fate-committed daughter cells become resistant to this defense mechanism both remain elusive.

Along a similar line of reasoning, it is plausible to predict that such a protective mechanism should also be activated when pervasive injuries or teratogenic insults damage the adherens junctions in the prenatal brain. For example, germinal matrix hemorrhage, the most common neurological disease of preterm infants as well as microcephaly-associated environmental teratogens, such as the Zika virus and fetal alcohol exposure are all known to impair cell-cell adhesion in the germinative ventricular zone leading to abnormal delamination and subsequent depletion of RGPC pools^15–18^. However, the molecular link between adherens junction damage and cell death induced by injury or by teratogenic insults remains unknown.

In the present study, we tested the hypothesis that a yet unidentified safeguarding mechanism determines distinct cell fate after normal delamination of daughter cells or abnormal detachment of progenitor cells. We demonstrate that adherens junction impairment induces aberrant RGPC delamination, ectopic accumulation and caspase-mediated apoptosis in the subventricular and ventricular zones, whereas caspase inhibition not only prevents the evoked cell death, but also rescues radial migration into the cortical plate. Moreover, abhydrolase domain containing 4 (ABHD4), a serine hydrolase with a yet undefined neurobiological function was identified as a necessary and sufficient mediator for the elimination of abnormally detached cells. Accordingly, healthy delaminated neuroblasts switch off ABHD4 in parallel with their neurogenic commitment. Notably, ABHD4 is not required for the canonical form of developmentally controlled programmed cell death in the embryonic neocortex indicating that its function is specifically associated with pathological insults. In agreement with this possibility, our findings elucidate that ABHD4 is also essential for prenatal alcohol exposure-induced apoptosis providing insights into the mechanisms underlying cell death associated with fetal alcohol syndrome.

## Results

### Abnormal RGPC delamination triggers developmental anoikis

N-cadherin (encoded by the *Cdh2* gene) is the major molecular component of the adherens junction belt along the ventricular wall in the developing mammalian brain^5^. To interfere with cadherin-based cell-cell adhesions, we carried out *in utero* electroporation of a dominant-negative version of N-cadherin (*ΔnCdh2-GFP*) into the lateral ventricles at embryonic day 14.5 (E14.5). We exploited this conditional and sparse adherens junction injury protocol instead of the global loss-of-function approaches to restrict the effects in time and space to a selected population of RGPCs and to avoid potential compensatory mechanisms that are more likely to come into play upon systems-level perturbations. Accordingly, *in utero* electroporation of *ΔnCdh2-GFP* caused a spatially limited destruction of adherens junctions around the transfected RGPCs (Fig. 1a-d; Supplementary Fig. 1a-f). STORM super-resolution imaging and confocal microscopy revealed the striking specificity of this experimental manipulation, because neither the basal endfeet at the pial surface nor the nanoscale architecture of radial scaffold processes of transfected RGPCs were impaired one day later (Supplementary Fig. 1g-r). As a result of the selective adherens junction disruption, the detached RGPCs delaminated from the ventricular zone and accumulated in the subventricular zone. These abnormally delaminated progenitor cells retained PAX6 transcription factor expression, a marker for proliferating RGPCs^19^ even outside of the ventricular germinative niche (Fig. 1e-i).

**Fig. 1.**
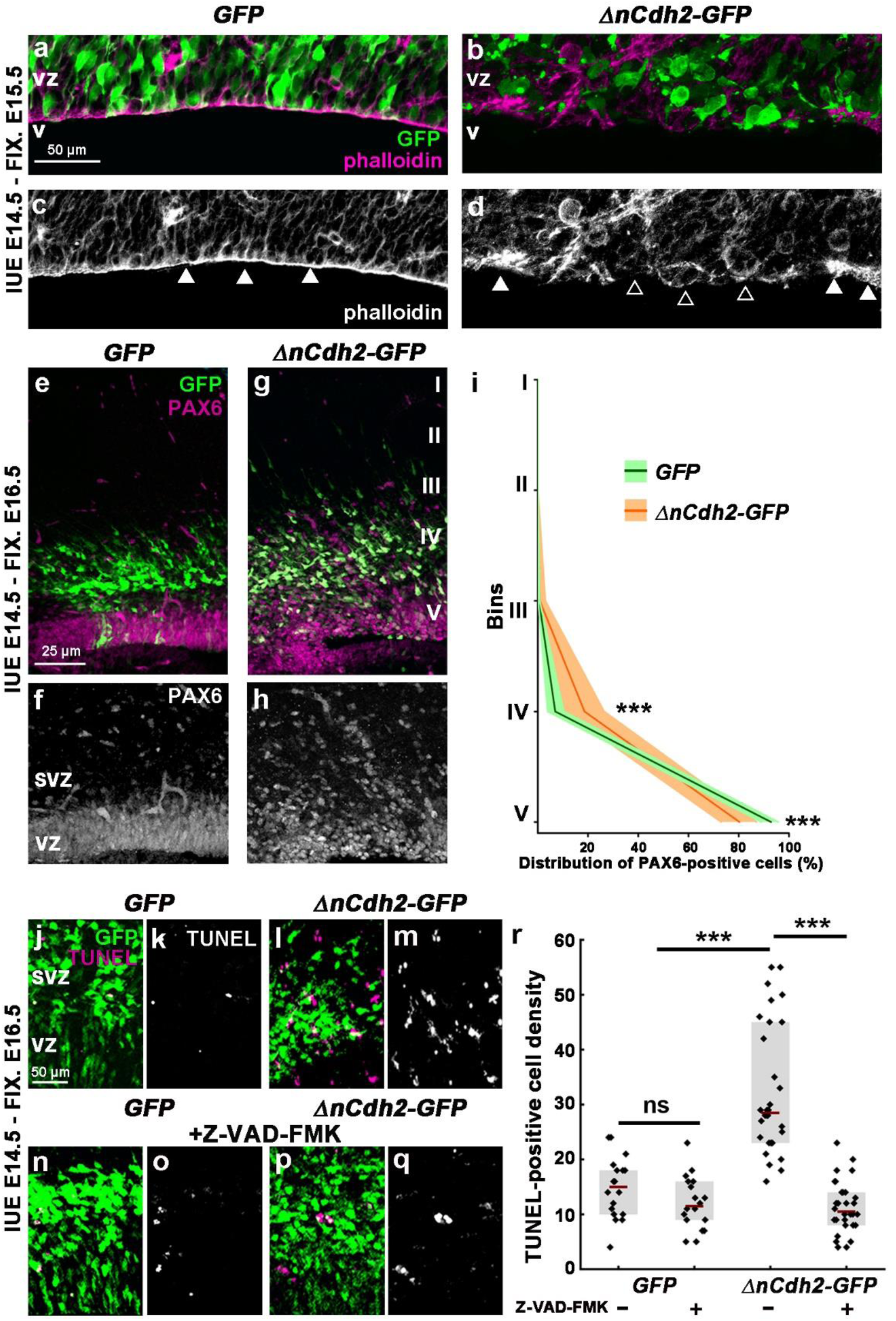
Adherens junction disruption induces apoptosis in the prenatal neocortex. **a-d**, High-power fluorescent micrographs of the ventricular zone (VZ) show the adherens junction belt (white arrowheads) around the ventricle (V). *ΔnCdh2-GFP-* (**b,d)**, but not control *GFP-in utero* electroporation (IUE) (**a,c)**, demolishes adherens junctions (open arrowheads). **e-h,** Breakdown of adherens junctions prompts delamination (**g-h**) and ectopic accumulation of PAX6-positive cells in the subventricular zone. **i,** Distribution of PAX6-positive cells in five equal bins (Roman numerals) (two-sided Mann-Whitney U test for all comparisons; 4^th^ bin \*\*\**P* = 0.0004 and 5^th^ bin \*\*\**P* = 0.0003; n = 13 sections from n = 3 animals per GFP treatment; n = 11 sections from n = 3 animals per *ΔnCdh2-GFP*-treatment). Data are shown as median (line) and interquartile range (transparent band). **j-m,** Confocal images demonstrate that *ΔnCdh2-GFP*- (**l,m)**, but not *GFP*-electroporation (**j,k)** triggers cell death. **n-q,** The pan-caspase inhibitor Z-VAD-FMK prevents cell death induced by *ΔnCdh2-GFP-*electroporation. **r,** Density of TUNEL-positive dead cells in the subventricular (SVZ) and ventricular zone (VZ) (Kruskal-Wallis test with post hoc Dunn’s Test; *\*\*\*P* < 0.0001; ns = not significant, *P ≈* 1; *n =* 18-18 sections from *n* = 3-3 animals per *GFP* and *GFP*+Z-VAD-FMK treatment; *n =* 30-30 sections from *n =* 4-4 animals per *ΔnCdh2-GFP* and *ΔnCdh2-GFP*+Z-VAD-FMK treatment). Graphs show raw data and median ± interquartile range.

In order to investigate the fate of these prematurely delaminated RGPCs, we next tested whether cell death is also induced by adherens junction disruption. Notably, *ΔnCdh2-GFP*-electroporation caused a strong increase in the density of dead cells when compared to *GFP*-electroporation in control animals. Administration of the pan-caspase inhibitor Z-VAD-FMK fully prevented the increase in cell death evoked by adherens junction disruption. In contrast, basal cell death levels remained unaffected demonstrating that abnormal delamination specifically triggers a caspase-dependent apoptotic process (Fig. 1j-r). Moreover, the ectopic accumulation of RGPCs in the subventricular zone was also rescued by Z-VAD-FMK-treatment and most of the surviving *ΔnCdh2-GFP*-electroporated cells could migrate to the cortical plate (Supplementary Fig. 2). These observations reinforce the idea that a specific cell death mechanism exists to eradicate prematurely delaminated progenitor cells in the developing neocortex, and this mechanism must be switched off in normally delaminating daughter cells to permit their normal migration to their functional destinations. We termed this process developmental anoikis, because it conceptually resembles the specific form of apoptosis of epithelial cells induced by the loss of cell anchorage^20, 21^. Anoikis has primarily been implicated as a protective mechanism in tumor biology, and pathologically detached tumor cells need to develop resistance to anoikis to become able to invade distant organs during metastasis formation^22^.

### Selective ABHD4 expression in RGPCs in the prenatal brain

The previous experiments led us to hypothesize that specific molecular players have evolved to mediate delamination error-induced cell death. These signaling components must be present in anchorage-dependent RGPCs to protect them from pathological insults, but their expression should be downregulated in migrating healthy neuroblasts. Therefore, we searched public single-cell RNA-Seq databases, and identified abhydrolase domain containing 4 (ABHD4), an endocannabinoid-metabolizing enzyme^23^ with a yet unknown *in vivo* function, as a potential molecular candidate. Notably, *Abhd4* mRNA was found to be highly expressed in putative RGPC pools in mouse and human embryonic cortical samples and cerebral organoids, whereas its level was below detection threshold in more committed neuronal progenitor cell populations and in adult cortical neuronal types^24–27^. Moreover, a target-agnostic *in vitro* shRNA library screen in immortalized prostate epithelial cells suggested that downregulation of *Abhd4* may potentially contribute to the resistance to anoikis^28^.

To experimentally determine whether the expression pattern of ABHD4 would suit its functional role in the developing neocortex, we first performed *in situ* hybridization on wild-type and littermate *Abhd4-*knockout embryonic brains. These experiments revealed that *Abhd4* mRNA expression was remarkably abundant in the germinative niches around all four brain ventricles, whereas it was absent in other regions and in control *Abhd4*-knockout embryonic brains. This spatially restricted expression pattern was consistent throughout prenatal development (Fig. 2a-e; Supplementary Fig. 3a-f), but *Abhd4* expression markedly decreased postnatally in parallel with the reduced number of proliferating progenitors in the subventricular and subgranular zones (Fig. 2f-h; Supplementary Fig. 3g-i), reaching undetectable levels in adults. Immunoblotting with a novel antibody raised against a conserved disordered motif of the ABHD4 protein further confirmed the presence of this serine hydrolase enzyme in the developing neocortex of wild-type, but not of *Abhd4-*knockout control embryos (Supplementary Fig. 3j,k).

**Fig. 2.**
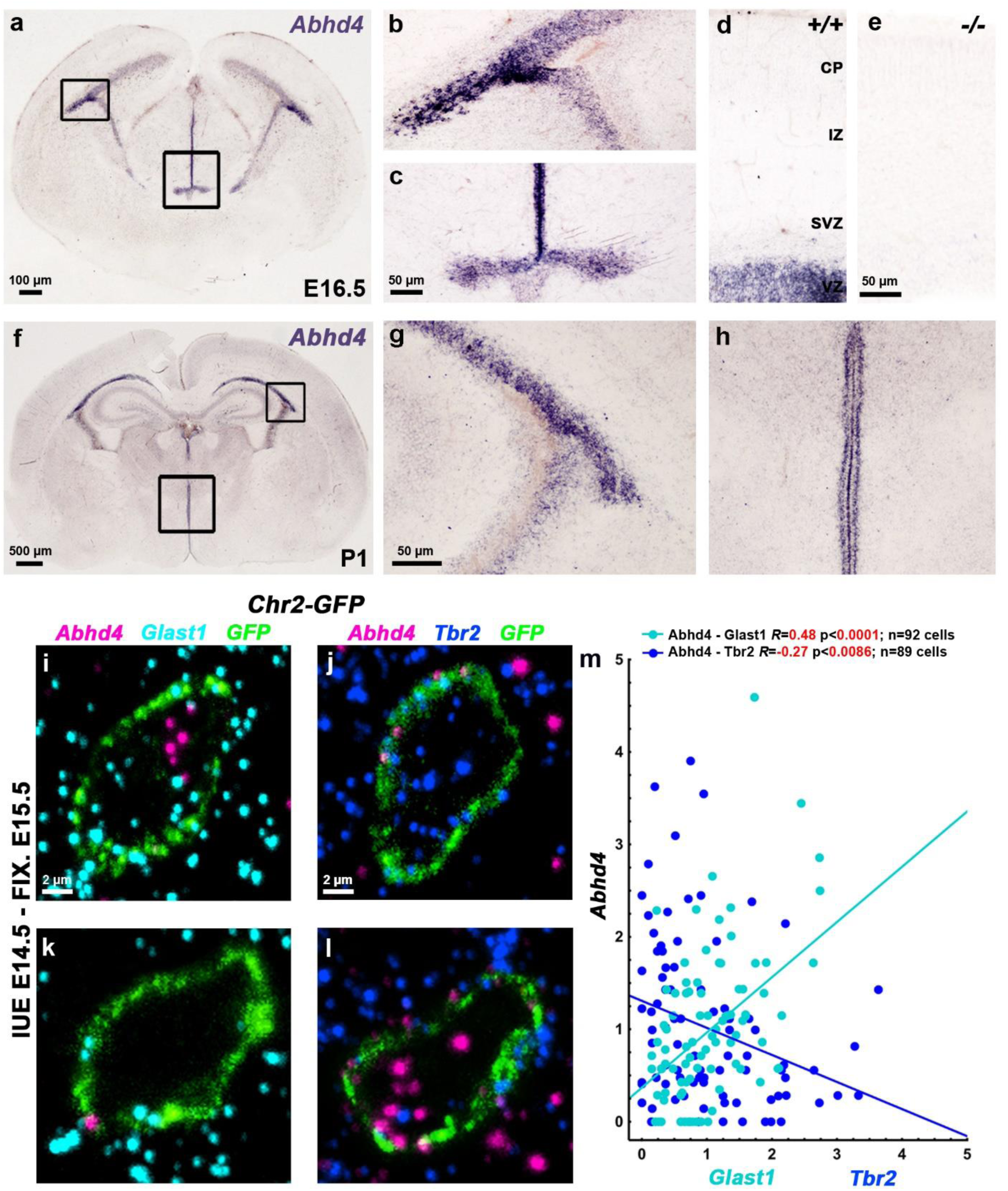
*Abhd4* mRNA is expressed by radial glia progenitor cells in the germinative niches. **a-h,** *Abhd4* mRNA is present exclusively in the ventricular zone along the lateral (**b,g**) and third ventricles (**c,h**) at both E16.5 (**a-d**) and P1 (**f-h**) wild-type (+/+) mice. The specificity of the *Abhd4* riboprobe is validated in *Abhd4-knockout* (-/-) animals (**e**). CP, cortical plate; IZ, intermediate zone; SVZ, subventricular zone; VZ, ventricular zone. **i-l,** High-power confocal imaging outlines the plasma membrane of *ChR2-GFP*-electroporated cells and delimits multi-color RNAscope analysis into single cells within the heterogeneous and densely packed cell layer of the ventricular zone. *Abhd4* mRNA typically colocalizes with the radial glia progenitor cell marker *Slc1a3* mRNA (encoding GLAST1 protein) (**i**), whereas other cells are often devoid of both markers (**k**), or either express *Eomes* (encoding TBR2 protein), a marker for committed daughter cells (**j**), or *Abhd4* alone (**l**). **m,** Correlation analysis of *Abhd4* mRNA levels with *Glast1* or *Tbr2* mRNA levels in single cells (Spearman’s rank correlation, *Abhd4/Glast1: R =* 0.4814, *P <* 0.0001; *n =* 92 cells from *n =* 4 mice; *Abhd4/Tbr2: R =* −0.241, *P =* 0.0221; *n =* 90 cells from *n =* 4 mice). The scatter plot shows data from individual cells normalized to the median value of the respective mRNA levels.

Although RGPCs represent the majority of cells in the germinative niches, it is important to note that fate-committed daughter cells that are undergoing delamination still populate the ventricular zone, where the high cellular abundance renders cell-specific quantitative mRNA analysis very difficult. In order to unequivocally identify the cell population expressing *Abhd4*, we developed a new approach for single cell-restricted, high-resolution visualization of the plasma membrane via *in utero* electroporation combined with multi-color *in situ* hybridization (see Online Methods). We found that *Abhd4* mRNA levels were positively correlated with *Glast1* expression (a marker of RGPCs^29^) and were inversely correlated with *Tbr2* expression (a marker of intermediate progenitor cells^30^) in daughter cells still residing within the ventricular zone (Fig. 2i-m). These findings together indicate that the temporal expression of *Abhd4* in RGPCs is tightly controlled and its rapid downregulation in fate-committed daughter cells starts immediately after their delamination.

### ABHD4 is not required for traditional RGPC functions, but it is sufficient to trigger apoptosis

The previous experiments demonstrated strict spatial and temporal restriction of *Abhd4* expression in RGPCs of the developing brain. RGPCs serve two major functions in the embryonic cerebral cortex: as progenitor cells^1–3^ and by providing a scaffold for postmitotic neuroblast migration^10^. By using *Abhd4*-knockout mice, we first tested the possibility that ABHD4 plays a role in RGPC proliferation. However, neither single-pulse bromodeoxyuridine-labeling of proliferating precursors (Fig. 3a-c), nor direct visualization of mitotic cells by phospho-histone H3-immunostaining (Supplementary Fig. 4a-c) revealed any quantitative differences between wild-type and *Abhd4-*knockout mice. This corresponds well with the lack of any changes in the number of Pax6-positive RGPCs (Fig. 3d-f) and in the structure of the adherens junction belt in the ventricular wall (Fig. 3g,h), together indicating an intact and functional neurogenic niche in *Abhd4-*knockout mice. Delamination and migration did not require ABHD4 either, because there was no change in the number and laminar distribution of TBR2-immunopositive intermediate progenitor cells^30^ in the subventricular zone and the density of TBR1-immunopositive postmitotic neurons^31^ was also similar in the cortical plate of the embryonic cerebral cortex (Supplementary Fig. 4d-i). Moreover, the laminar distribution of pyramidal cells, GABAergic interneurons and astrocytes was also unaltered in the adult neocortex (Supplementary Fig. 5). Finally, the nanoarchitecture of the radial glia processes also remained intact in the absence of ABHD4 (Fig. 3i-p). Together, these results demonstrate that despite its high expression in RGPCs, ABHD4 is not necessary for the two major developmental functions of these progenitor cells.

**Fig. 3.**
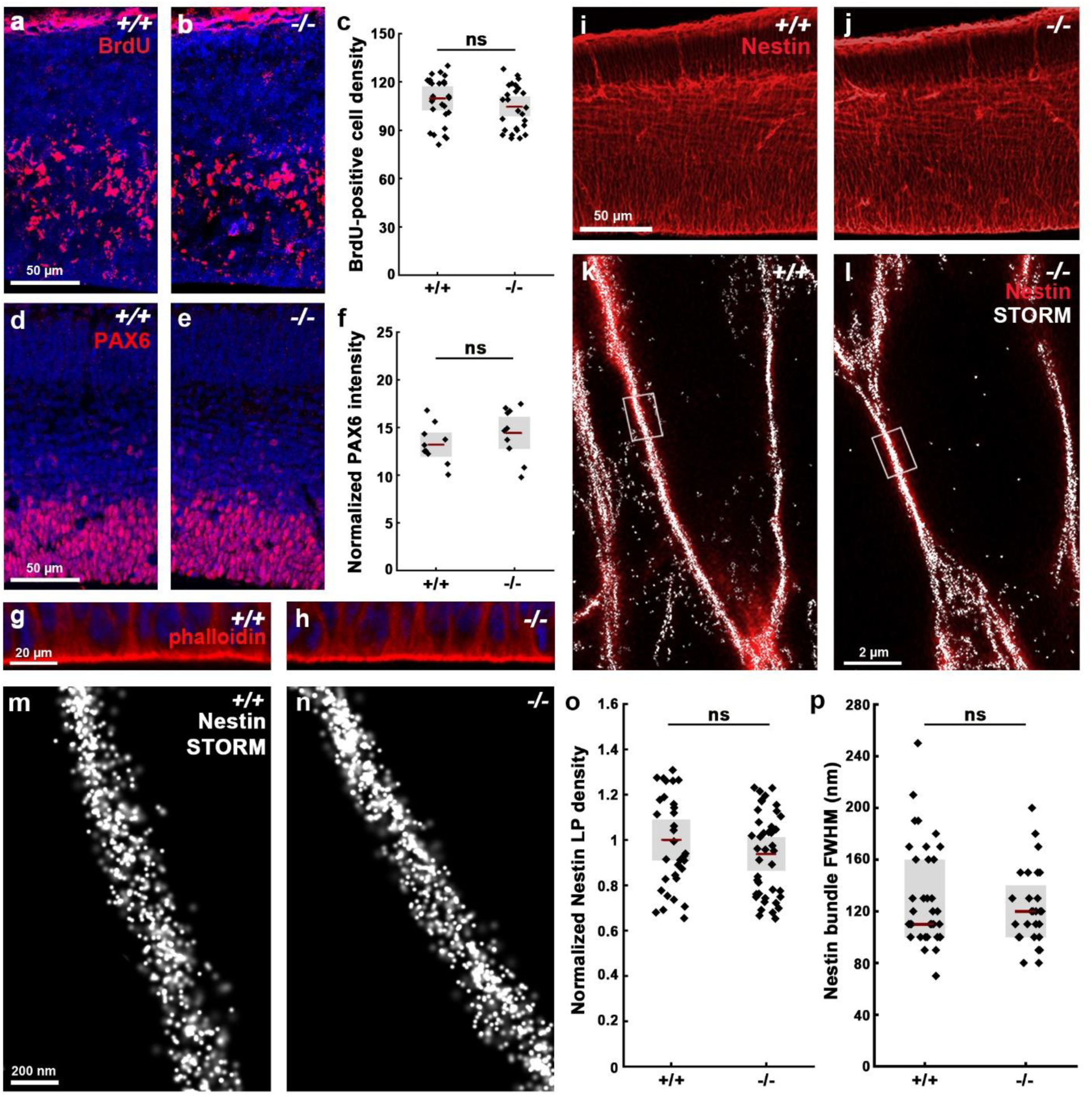
ABHD4 is not required for classical radial glia progenitor cell functions. **a,b,** Coronal sections of the embryonic cortex at E.14.5 show replicating cells that incorporated bromodeoxyuridine (BrdU) during S phase of the cell cycle are shown. **c,** Quantification of BrdU-positive cell density (two-sided Student’s unpaired t-test, *P =* 0.323; *n =* 38 sections from *n =* 6 animals per wild-type (+/+) mice, *n =* 29 sections from *n =* 4 animals per *Abhd4-*knockout (-/-) mice). **d,e,** Confocal images and **(f)** quantification of PAX6-positive radial glia progenitor cells in the ventricular zone. (two-sided Student’s unpaired t-test, *P =* 0.2598; *n =* 10 sections from *n =* 4 animals per wild-type (+/+) mice, *n =* 10 sections from *n =* 4 animals per *Abhd4-* knockout (-/-) mice). **g,h,** Identical pattern of phalloidin-labeling of the adherens junction belt in the ventricular zone (VZ) of wild-type (+/+) and *Abhd4-*knockout (-/-) mice at E15.5. **i,j** Low-power confocal images show similar organization of nestin-positive radial processes in both genotypes at E15.5. **k,l,** Correlated confocal and STORM super-resolution microscopy images from both genotypes. **m,n,** STORM super-resolution images reveal that the nanoarchitecture of nestin intermediate filament bundles at the core of the radial glia processes remains intact in the absence of ABHD4. **o,** Density of localization points (LPs) representing nestin (two-sided Student’s unpaired t-test, *P =* 0.297; *n =* 39 segments from *n =* 3 animals per wild-type (+/+) mice, *n =* 57 segments from *n =* 3 animals per *Abhd4-*knockout (-/-) mice). Graphs show raw data and mean ± 2 x standard error. **p,** Quantification of full-width-at-half-maximum of nestin filament bundles (two-sided Mann-Whitney U test, *P =* 0.684; *n =* 28 segments from *n =* 3 animals per wild-type (+/+) mice, *n =* 37 segments from *n =* 3 animals per *Abhd4-*knockout (-/-) mice). Graphs show raw data and median ± interquartile range.

These unexpected findings, together with the spatially and temporally restricted expression of *Abhd4* in RGPCs, raise the intriguing possibility that this serine hydrolase has evolved to fulfill a hitherto undefined, but conserved physiological function that is important in the cell biology of neurogenesis. In agreement with this hypothesis, a phylogenetic analysis revealed that ABHD4 protein orthologues exhibit substantial homology that is especially high around the catalytic serine residue (S159) and the consensus “nucleophile elbow” sequence “GXSXG”, suggesting that the enzymatic function of ABHD4 is evolutionarily conserved (Supplementary Fig. 6).

Because *Abhd4* expression is rapidly downregulated in delaminated daughter cells right after their fate-commitment, we reasoned that counteracting this process by ectopic *Abhd4* expression in these cells could shed light on the conserved function of ABHD4 in the developing brain. To test this idea, we *in utero* electroporated either *GFP* alone, or wild-type *Abhd4-GFP*, or a catalytically inactive form of *Abhd4-GFP* (in which the catalytic serine residue that is required for hydrolase activity was mutated to glycine, S159G mutant) at E14.5. Interestingly, ABHD4, but not the inactive, hydrolase-dead form of ABHD4 caused a striking migration arrest within the subventricular zone (Fig. 4a-d). Moreover, *Abhd4-GFP*-transfected cells lost the characteristic bipolar morphology of migrating neuroblasts and became shrunken, rounded and had no leading processes (Fig. 4e-h). These morphological changes are hallmarks of imminent cell death. Indeed, we observed that ABHD4 alone, but not its inactive form could trigger a robust (295%) increase in caspase-dependent apoptosis (Fig. 4i-o) when compared to *GFP-* electroporated controls, indicating that the enzymatic activity of ABHD4 is necessary and sufficient to elicit apoptosis in the delaminated cells.

**Fig. 4.**
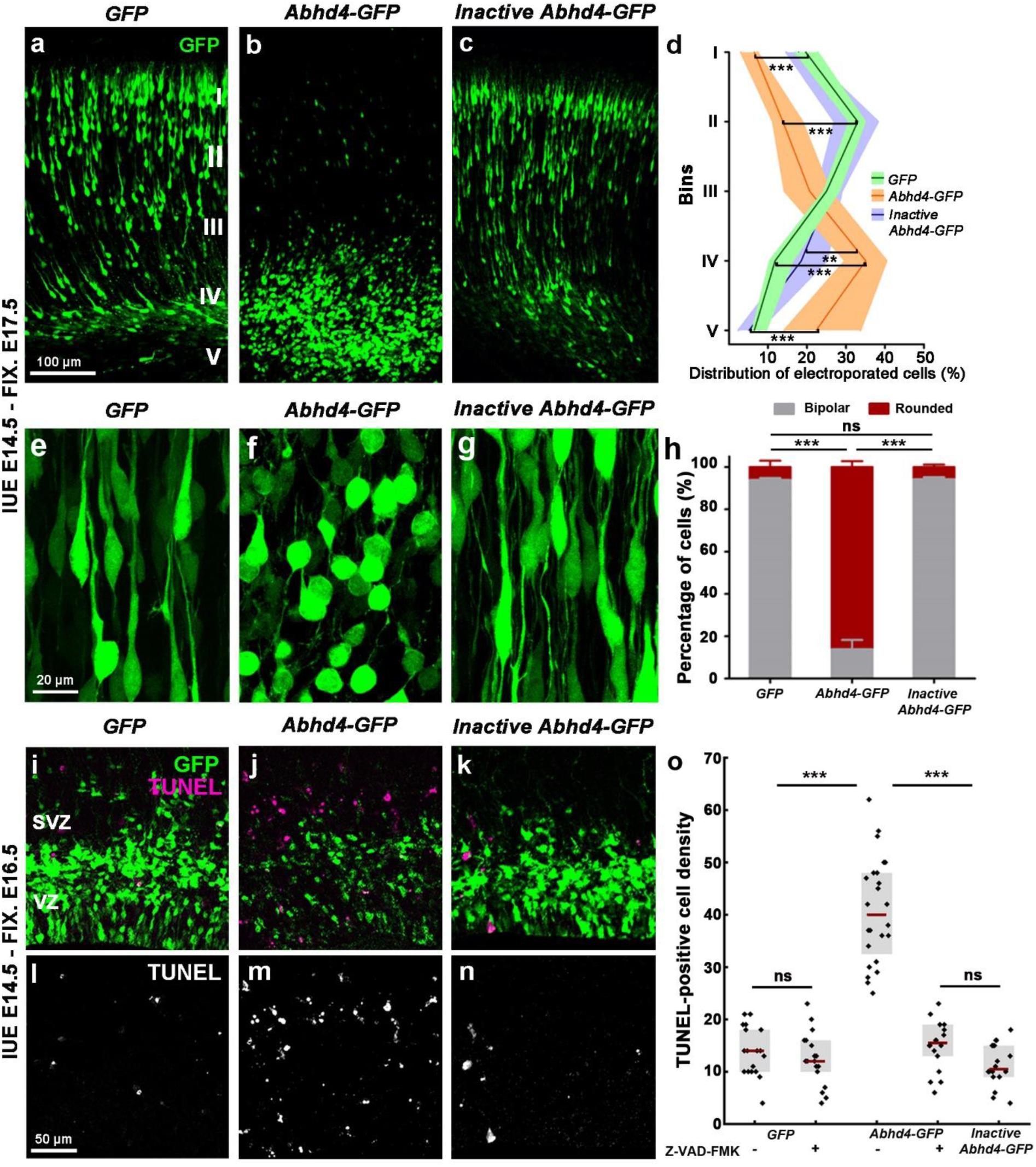
ABHD4 is sufficient to trigger migration arrest and apoptosis. **a-c,** Cells *in utero* electroporated (IUE) with *GFP-* (**a**), or with an *Abhd4-GFP*-construct encoding an enzymatically inactive form of ABHD4 (**c**) migrate into the cortical plate, whereas *Abhd4-GFP*-electroporation causes radial migration defect (**b**). **d,** Laminar distribution of electroporated cells in five equally-sized bins (Roman numerals) (Kruskal-Wallis test with post hoc Dunn’s test, 1^st^,2^nd^,4^th^,5^th^ bins for all comparisons: \*\*\**P <* 0.0001, except 4^th^ bin (*Inactive Abhd4-GFP* vs *Abhd4-GFP*): \*\**P =* 0.006; *n =* 19 sections from *n =* 4 animals per *GFP*-electroporation; *n =* 18 sections from *n = 4* animals per *Abhd4-GFP*-electroporation; *n =* 12 sections from *n =* 3 animals per Inactive *Abhd4-GFP*-electroporation). Data are shown as median (line) and interquartile range (transparent band in the same color). **e-g,** High-power images show bipolar migrating neurons (**e, g**), that become rounded upon *Abhd4-GFP*-electroporation (**f**). **h,** Quantification of cell morphology (Kruskal-Wallis test with post hoc Dunn’s test, \*\*\**P <* 0.0001; ns = not significant, *P ≈* 1; *n =* 15-15 sections from *n* = 3-3 animals per treatment). Graphs show median ± interquartile range. **i-n,** Representative images show large density of TUNEL-positive cells upon *Abhd4-GFP* electroporation in the subventricular (SVZ) and ventricular (VZ) zone (**j,m**), that is not replicated by *GFP-* (**i,l**), or by inactive *Abhd4-GFP*-electroporation (**k,n**). **o,** Quantification of TUNEL-positive cell density (Kruskal-Wallis test with post hoc Dunn’s test, \*\*\**P <* 0.0001; ns = not significant, *P ≈* 1 except per *Abhd4-GFP*-electroporation with Z-VAD-FMK treatment vs *Inactive Abhd4-GFP P =* 0.607; *n =* 18-18 sections from *n =* 3-3 animals per *GFP*-electroporation with and without Z-VAD-FMK treatment and per Inactive *Abhd4-GFP*-electroporation; *n =* 24 sections from *n =* 4 animals per *Abhd4-GFP*-electroporation; *n =* 18 sections from *n =* 4 animals per *Abhd4-GFP*-electroporation with Z-VAD-FMK treatment). Graphs show raw data and median ± interquartile range.

To determine how powerfully ABHD4 drives cell death even without a native tissue context (e.g. in the absence of additional pro-apoptotic signals), we next examined Human Embryonic Kidney-293 cells, which are relatively resistant to pro-apoptotic stimuli due to their deregulated pRB/p53 pathway and also lack endogenous *Abhd4* expression^32^. Correlated confocal and STORM super-resolution imaging revealed that ABHD4, but not its inactive form caused a substantial loss of TOM20-positive mitochondria (Supplementary Fig. 8a-f,g) and a concomitant rise in extra-mitochondrial cytochrome c levels, signs of early stage apoptosis (Supplementary Fig. 8e,f,h-j). Moreover, ABHD4 also elicited a large increase in the number of cleaved caspase 3-immunopositive cells exhibiting nuclear fragmentation and condensation, features of late stage apoptosis (Supplementary Fig. 8k-q). In contrast, the adjacent untransfected cells remained unaffected (Supplementary Fig. 8r). These results indicate that ABHD4 is capable of triggering an apoptotic program cell-autonomously, and even in cell types with a higher apoptotic threshold.

### ABHD4 is not required for developmentally controlled programmed cell death, but it is necessary for developmental anoikis

It is well-established that cell type-specific consecutive waves of developmentally controlled programmed cell death delete a substantial proportion of neurons during cortical development^33–35^. Considering the pro-apoptotic activity of ABHD4, we therefore investigated the basal level of cell death in the developing neocortex. Notably, there was no difference in the density of dead cells in the subventricular and ventricular zones of wild-type and *Abhd4*-knockout mice at embryonic day 16.5 (Fig. 5a-e). Developmentally controlled programmed cell death of glutamatergic cells has another peak early postnatally^34^, but there were no differences in cell death between the genotypes even at postnatal day 3 (Supplementary Fig. 8).

**Fig. 5.**
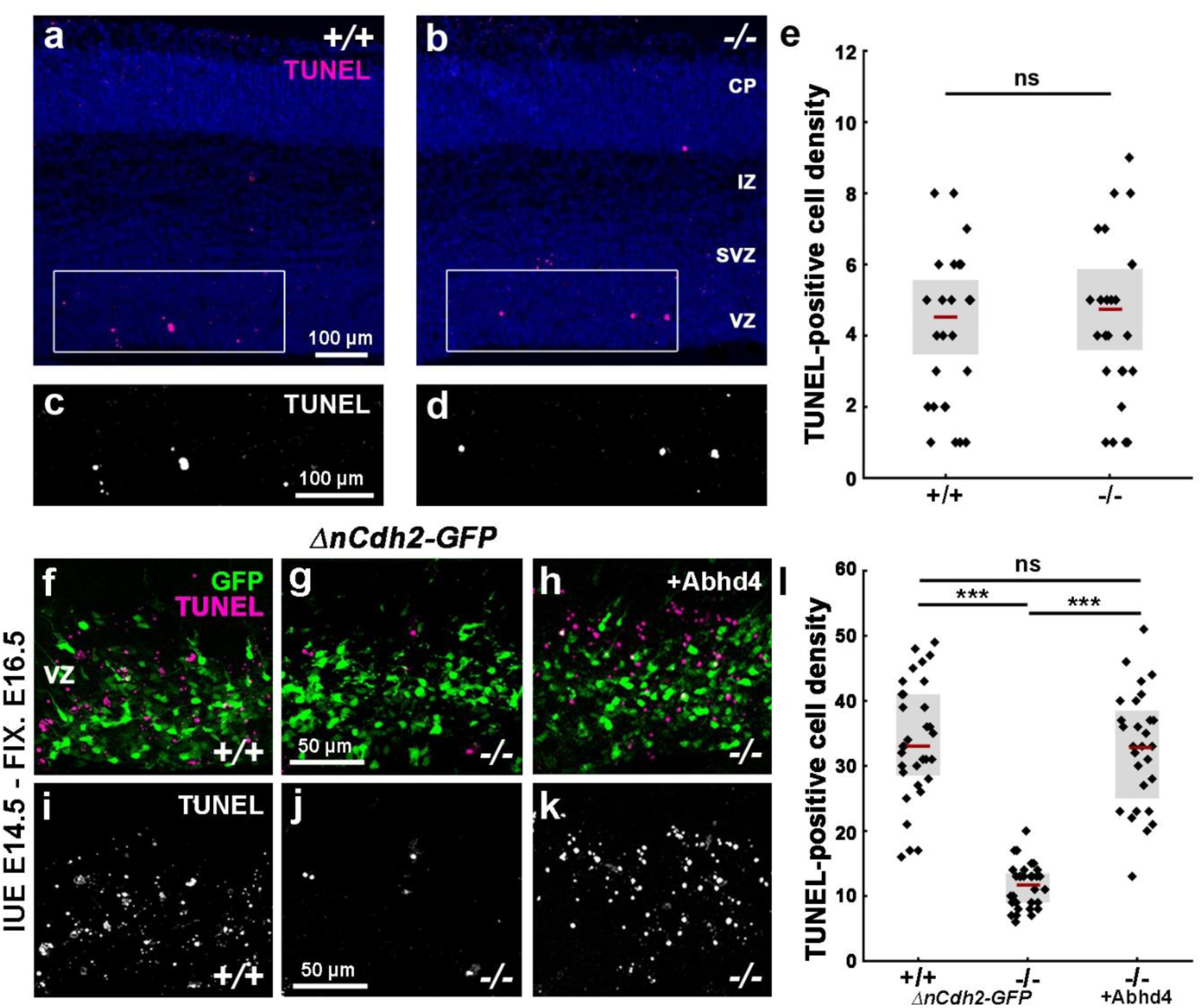
ABHD4 mediates apoptosis induced by the loss of adherens junctions, but it is not required for developmentally controlled programmed cell death. **a-d,** Basal level of cell death is similar in coronal sections of the embryonic neocortex of wild-type (+/+) and *Abhd4-*knockout (-/-) mice at E16.5. CP, cortical plate; IZ, intermediate zone; SVZ, subventricular zone; VZ, ventricular zone. **e,** Quantification of TUNEL-positive cell density in the VZ/SVZ (two-sided Student’s unpaired t-test, *P =* 0.6964; *n =* 30 sample from *n =* 5 animals from both genotypes). **f,g,i,j,** *ΔnCdh2-GFP in utero* electroporation (IUE) triggers cell death in wild-type (**f,i**), but not in *Abhd4*-knockout mice (**g-j**) in the subventricular (SVZ) and ventricular (VZ) zones. **h,k,** Re-expression of *Abhd4-GFP* in *Abhd4*-knockout mice is sufficient to rescue impaired *ΔnCdh2-GFP*-induced cell death. **l,** Quantification of TUNEL-positive cell density (Kruskal-Wallis test with post hoc Dunn’s test, \*\*\**P <* 0.0001; ns = not significant, *P* ≈ 1; *n =* 32-32 sections from *n =* 4-4 animals per genotypes, *n =* 28 sections from *n =* 4 animals per *Abhd4*-re-expression treatment). Graphs show raw data and median ± interquartile range.

However, physiologically controlled forms of programmed cell death or apoptosis induced by pathological insults do not necessarily share the same molecular mechanisms. Because our results have revealed that both adherens junction disruption-induced abnormal delamination and ectopic ABHD4 enzymatic activity in normally delaminated cells cause similar caspase-dependent cell death in the embryonic neocortex, we next tested the possibility that ABHD4 serves as a specific molecular link between abnormal delamination and subsequent cell death. To this end, *ΔnCdh2-GFP* was electroporated into the lateral ventricles of wild-type and littermate *Abhd4-*knockout embryos. Sparse adherens junction disruption initiated a marked increase in the density of dead cells in wild-type mice that was completely absent in *Abhd4-*knockout mice (Fig. 5f,g,i,j,l). Importantly, this increase in cell death could be fully rescued by ABHD4 re-expression in *Abhd4-*knockout mice (Fig. 5h,k,l). These findings demonstrate that ABHD4 is not required for developmentally controlled programmed cell death, but it is essential for developmental anoikis, a unique form of cell death triggered by the damage of cadherin-based adherens junctions in the prenatal brain.

### Maternal alcohol exposure triggers ABHD4-dependent cell death in the embryonic neocortex

The most common preventable teratogenic insult in humans is alcohol. Fetal alcohol spectrum disorder has a remarkably high prevalence and it is the foremost non-genetic cause of intellectual disabilities. Prenatal ethanol exposure can cause several brain malformations including cortical dysplasia/heterotopias and microcephaly, all indicating impaired neurogenesis^16^. Importantly, ethanol is known to induce disruption of the adherens junction signaling machinery, to reduce the RGPC pool and to increase cell death, including anoikis^15, 16, 36, 37^. Therefore, we hypothesized that ABHD4 may play a role in cell death associated with prenatal alcohol exposure. We tested this idea by applying two independent ethanol administration protocols to pregnant mice. Both a 3-day-long subchronic ethanol administration regime and a single dose of acute ethanol administration resulted in a substantial (3-6 fold) increase in cell death and led to the scattered distribution of dead cells throughout the ventricular and subventricular zones in wild-type embryos (Fig. 6). In striking contrast, maternal ethanol administration failed to induce a similar increase in cell death in littermate *Abhd4-*knockout embryos (Fig. 6d,h,i,m,q,r). These results strongly suggest that ABHD4 is also indispensable for ethanol-induced cell death in the embryonic brain.

**Fig. 6.**
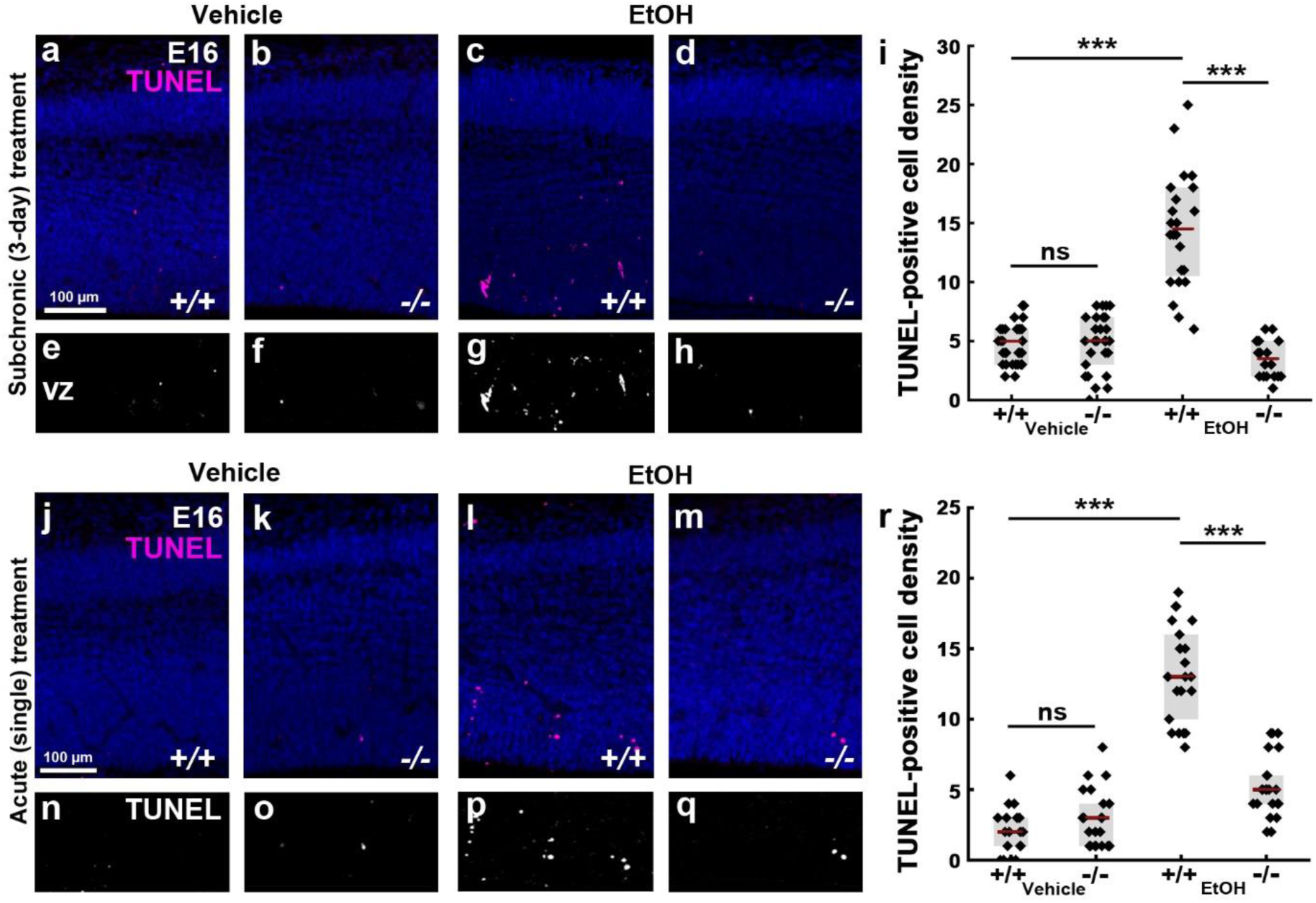
ABHD4 is necessary for cell death caused by fetal alcohol exposure. **a-h**, Compared to vehicle-treated mice (**a,b,e,f**), increased number of TUNEL-positive dead cells is seen in ethanol (EtOH)-treated wild-type (+/+), (**c,g**), but not in *Abhd4-*knockout (-/-) mice (**d,h**) in the subventricular (SVZ) and ventricular (VZ) zones of embryos derived from dams undergoing subchronic alcohol treatment. **i,** Quantification of the density of TUNEL-positive cells in the subchronic model (Kruskal-Wallis test with post hoc Dunn’s test, \*\*\**P <* 0.0001; ns = not significant, *P ≈* 1, *n =* 31 sections from *n =* 5 animals per wild-type, vehicle-treated mice, *n =* 27 sections from *n =* 4 animals per *Abhd4-*knockout, vehicle-treated mice, *n =* 24 sections from *n =* 4 animals per wild-type, ethanol-treated mice, *n =* 18 sections from *n =* 3 animals per *Abhd4-*knockout, ethanol-treated mice). **j-q,** A single maternal exposure to alcohol evokes cell death in the developing neocortex of wild-type (**l,p**), but not in *Abhd4-*knockout embryos (**m,q**). **r,** Quantification of cell death in the acute alcohol model (Kruskal-Wallis test with post hoc Dunn’s test, \*\*\**P <* 0.0001; ns = not significant, *P ≈* 1; *n =* 18 sections from *n =* 3 animals per wild-type, vehicle-treated mice, *n =* 21 sections from *n =* 3 animals per *Abhd4-*knockout, vehicle-treated mice, *n =* 22 sections from *n =* 3 animals per wild-type, ethanol-treated mice, *n =* 23 sections from *n =* 3 animals per *Abhd4-*knockout, ethanol-treated mice). Graphs show raw data and median ± interquartile range (**i, r**).

## Discussion

In the present study, we provide evidence for the existence of developmental anoikis, a distinctive mechanism for cell death in the prenatal brain. Our findings also identify ABHD4, an enzyme that has a vital function in this phenomenon by preventing the survival of misplaced cells that are produced by delamination errors or damaged by alcohol exposure (Supplementary Fig. 9). Various environmental stressors are known to affect the fetal brain and considering the immense number of cell division events and subsequent delamination steps (>10^11^), the frequency of teratogenic and spontaneous delamination errors is likely to be considerably higher than the observed incidence of congenital brain anomalies (∼0.25%, https://eu-rd-platform.jrc.ec.europa.eu/eurocat/eurocat-data/prevalence). Thus, it is conceivable to postulate the pivotal neurological importance of ABHD4-dependent developmental anoikis that eradicates those abnormally delaminated cells that may represent a risk for future brain malformations, such as dysplasias, heterotopias or even brain tumors.

It is important to emphasize that RGPCs have a neuroepithelial origin. These proliferating cells maintain most of the epithelial characteristics including the apico-basal polarity via specific molecular anchors to the extracellular matrix and to each other^6^. Right after cell division, the delaminating non-RGPC-fated daughter cell dismantles its cell-cell connections, switches its transcriptional profile and undergoes a characteristic cytoskeletal reorganization to initiate migration. It is increasingly appreciated that several molecular components and the cellular processes that underlie daughter cell delamination are very similar in the so-called epithelial-mesenchymal transition phenomenon^8, 38^. Our study further adds to this idea, because the ABHD4-dependent cell death mechanism described here is conceptually analogous to anoikis, a detachment-induced form of apoptosis that has long been considered as the central protective mechanism in epithelial-mesenchymal transition to prevent the proliferation and migration of abnormally delaminated epithelial cells^20, 21, 39, 40^. Although the term has originally been coined for intrinsic apoptosis induced by the loss of the extracellular matrix-dependent anchorage of epithelial cells^20, 21^, we propose to call this process developmental anoikis, because of its conceptual similarity and its functional importance during development. Traditionally, anoikis has primarily been implicated as a protective mechanism in tumor biology, and obtaining resistance to anoikis is a critical step for cancer cells for tumor invasion and metastasis^21, 22, 40^. Thus, the results that ABHD4 is a pro-apoptotic molecule whose expression rapidly disappears in daughter neuroblasts and coincidentally, that ABHD4 down-regulation in immortalized prostate epithelial cells prevents cell death^28^ together raise the intriguing possibility that a similar mechanism underlies anoikis resistance in healthy migrating neurons and in metastatic cancer cells. From a clinically relevant perspective, another interesting related finding is that ethanol-induced cell death also requires ABHD4 in the embryonic neocortex. Because alcohol turns on the epithelial-mesenchymal transition program that is thought to contribute to the metastatic aggressiveness of epithelial cancers associated with chronic alcohol consumption^41^, ABHD4-dependent cell death may play an important role in the attenuation of tumor progression. Notably, data from expression arrays indicate that the *Abhd4* gene is an important downstream target of the tumor suppressor protein p53 in cancer cells at the transcriptional level ^42, 43^. Thus, it remains to be investigated in future studies whether and how p53 and other related transcriptional regulators contribute to the tightly controlled spatial and temporal expression of *Abhd4* mRNA in the developing brain and under pathological conditions.

At the biochemical level, ABHD4 is involved in N-acyl-ethanolamine and related N-acyl-phospholipid metabolism including the production of the endocannabinoid molecule anandamide^23, 44^. Anandamide has various physiological functions including the regulation of synaptic plasticity and apoptosis^45^. Interestingly, NAPE-PLD, another anandamide-synthesizing enzyme is primarily concentrated in presynaptic axon terminals^46^ and its enzymatic activity only appears in the postnatal brain^47^. In contrast, our findings demonstrated that ABHD4 is present in radial glia progenitor cells in the prenatal brain, but it disappears postnatally. Together with the biological function of ABHD4 in the elimination of abnormally detached cells by triggering apoptosis, it is plausible to propose that the multiple enzymatic routes of anandamide synthesis may have evolved to maintain a division of labor of anandamide signaling in time and space. It is likely that in the context of these two very distinct biological phenomena, synaptic plasticity and apoptosis, anandamide signaling requires different regulatory mechanisms, which can be achieved by independently controlling gene expression and enzymatic activity of the different molecular components of the endocannabinoid system.

Besides the potential pro-apoptotic role of the signaling molecule anandamide produced by ABHD4 activity^45^, one must also consider the function of N-acyl-phosphatidylethanolamines (NAPEs), which are the direct substrates of ABHD4^44^. Early biophysical studies reported a stabilizing effect of NAPEs on the integrity of biological membranes and increasing the NAPE content of liposomes inhibits their leakage^48, 49^. Thus, ABHD4, as a NAPE lipase may also impair intracellular membrane compartments by hydrolyzing NAPEs. Interestingly, anandamide triggers apoptosis via inducing endoplasmic reticulum stress^50^. It is tempting to speculate that the synergistic upstream and downstream legs of ABHD4 activity in combination would represent a very efficient and parsimonious mechanism that renders damaged cells more susceptible to additional pro-apoptotic processes. In light of the well-described pro-apoptotic role of phosphatidylserine however, the potential significance of the N-acyl-phosphatidylserines (NAPSs), another substrate class of ABHD4^44^, is also worthy of consideration in future studies.

Elucidation of the unique biological function of ABHD4 in the embryonic cerebral cortex will not only help to identify the specific lipid mediators and promote the discovery of additional molecular players cooperating with ABHD4, but will also pave the way to delineate the precise biochemical cascades and biophysical processes of developmental anoikis. This is important, because the present findings indicate that ABHD4-mediated developmental anoikis is a safeguarding mechanism that protects the fetal brain from the effects of delamination errors induced spontaneously or by teratogenic insults. Thus, a better understanding of this phenomenon will also help to gain insights into the neurodevelopmental mechanisms underlying congenital brain anomalies and the related neurological and cognitive deficits.

## Online Methods

### Animals

All experiments were approved by the Hungarian Committee of the Scientific Ethics of Animal Research (license numbers: XIV-1-001/2332-4/2012 and PE/EA/354-5/2018), and were carried out according to the Hungarian Act of Animal Care and Experimentation (1998, XXVIII, Section 243/1998), in accordance with the European Communities Council Directive of 24 November 1986 (86-609-EEC; Section 243/1998). Mice were kept under approved laboratory conditions and all efforts were made to minimize pain and to reduce the number of animals used. Both male and female mice were used throughout the study. C57BL/6 and CD-1 mouse lines were obtained from Charles River Laboratories. Mice bearing a disruption in the *Abhd4* gene were generated from the 129S6/SvEvTac strain and were backcrossed into the C57BL/6 background for more than 10 generations prior to present experiments. The line has been validated in an earlier study^44^.

### DNA constructs and cloning protocols

Mouse N-cadherin (*Cdh2*) was cloned via RT-PCR using E16.5 brain cDNA as template with long-template PCR mix according to manufacturer’s instruction (Roche). The dominant-negative version of N-cadherin (*ΔnCdh2*) was created by OliI-HindIII digestion, followed by Klenow fill-in of HindIII-overhang and religation of the plasmid. The final construct was validated by sequencing and encoded a truncated N-cadherin molecule that contains the signal sequence followed by the transmembrane and intracellular domains but lacks the extracellular domain. The analogous mutant form of N-cadherin has been shown to interfere with cell adhesion in cellular assays derived from clawed frog and chicken^51–53^.

Mouse *Abhd4* was also amplified by RT-PCR as described above. The amplicon was cloned into pGEMTEasy plasmid (Promega) and sequenced. The construct encoding the inactive *Abhd4* form was created by site-directed mutagenesis of the catalytic serine residue 159 to glycine via PfuI PCR and DpnI digestion of the template plasmid, followed by transformation and clone identification by sequencing. The *ΔnCdh2*, *Abhd4* and inactive *Abhd4* constructs were subcloned from pGEMT into the pCAGIG mammalian expression vector. The pCAGIG plasmid was a gift from Connie Cepko (Addgene #11159)^54^.

To generate riboprobes for chromogenic in situ hybridization, a shorter 418bp-long fragment of the *Abhd4* gene and a 406 bp-long fragment of *Slc1a3* gene (encoding the astrocyte marker protein GLAST1) were amplified by RT-PCR and cloned into pGEMTEasy vector. The riboprobes encoded by the *Slc17a7* and *Gad1* genes that were used to visualize VGLUT1-positive pyramidal neurons and GAD67-positive interneurons in the neocortex, respectively, were made as reported before^55^.

The following primers were used for cloning: *Cdh2* (896 bp): forward primer 5’-ATGTGCCGGATAGCGGGAGCGC, reverse primer 5’-TCAGTCGTCACCACCGCCGTACATG; *Abhd4* (1068 bp): forward primer 5’-ATGGGCTGGCTCAGCTCGACCCG, reverse primer 5’-TCAGTCAACTGAGTTGCAGATCTCT; Inactive *Abhd4* (1068 bp): forward primer 5’-CCGTGCACCTCCAACCTGGGTCAGGGCTGTGGCATCTGTCC, reverse primer 5’-GGACAGATGCCACAGCCCTGACCCAGGTTGGAGGTGCACGG; short *Abhd4* (418 bp): forward primer 5’-CGGCAGGGCTTGTTTACTAT, reverse primer 5’-TCTCCCGCCATGTC TCTATT; *Slc1a3* (406 bp): forward primer 5’-TAGGGGCAGGCTGTGTGTGGCTCAC, reverse primer 5’-TCGTTTCTTTGGGGGCTGATTAGGGAC.

### In utero electroporation, Z-VAD-FMK and BrdU injections

Timed-pregnant female mice were anesthetized with avertin (1.25% v/v, Sigma), and uterine horns were exposed. The bicistronic construct harbouring the gene-of-interest and an internal ribosome entry site (IRES)-GFP cassette in the pCAGIG expression vector (1-2 μg/μl at 1 μl volume) in endotoxin-free water containing Fast Green (1:10000, Roth) was electroporated into the lateral ventricle of the embryo via a glass capillary at embryonic day 14.5. Electroporation was performed with tweezer electrodes (5 pulses of 40 V for 50 milliseconds at 950 millisecond intervals) using an SP-3c electroporator (Supertech). After electroporation, the uterine horns were returned into the abdominal cavity, the wall and skin were sutured and embryos were allowed to continue their normal development. To inhibit caspase activity, the pan-caspase inhibitor Z-VAD-FMK (5 μM, BD Biosciences) was injected in a similar manner as above. To identify proliferating cells, BrdU in 0.9% saline (200 mg/kg, Sigma) was intraperitoneally injected into pregnant dams at E14.5. The embryos were collected two hours later and their brains were fixed with 4% paraformaldehyde (PFA).

### Chromogenic and cell-specific RNAscope fluorescent *in situ* hybridization

Embryonic brains (E14.5, E16.5, E18.5) and early postnatal (P1, P3) brains were removed from the skull and immersion-fixed with 4% PFA overnight. Older postnatal (P10) and adult (P60) wild-type and littermate *Abhd4*-knockout mice were transcardially perfused with 4% PFA and their brains were postfixed for 3 hours at 4°C. After fixation, 50-μm-thick free-floating sections were cut with VT-1200S Vibratome (Leica). Chromogenic *in situ* hybridization using digoxygenin-labeled antisense riboprobes was performed as described earlier^56^.

The very high cellular abundance in the ventricular zone renders quantitative measurement of mRNA molecules in individual cells very difficult by conventional *in situ* hybridization approaches. In order to visualize the plasma membrane of a sparse population of cells in the ventricular zone by *in utero* electroporating *ChR2-GFP* into the lateral ventricles of embryos at E14.5. The AAV-CAG-ChR2-GFP plasmid was a gift from Edward Boyden (Addgene #26929)^57^. One day after electroporation, the brains of embryos were removed and frozen on isopentane chilled with dry ice. The brain tissue was equilibrated in the cryostat for 2-3 hours at −20°C, then 20-μm-thick cryosections were collected on Superfrost Ultra Plus glass slides (ThermoFisher) and were held in the cryostat until finishing the sectioning. Sections were fixed with 10% ice-cold PFA for 30 minutes at 4°C. After fixation, cryosections were washed with PBS and dehydrated with a series of alcohol solutions in ascending concentrations of 50% ethanol, 75% ethanol and absolute ethanol each for 5 minutes, then slides were transferred to the last jar containing absolute ethanol and were incubated overnight at −20°C. Next day, the RNAscope Multiplex Fluorescent Detection assay was performed based on manufacturer’s protocol (Advanced Cell Diagnostics). The Protease IV treatment was shortened to 10 minutes to preserve enough GFP protein in the plasma membrane for sharp contour of individual cells in the ventricular zone. The *Abhd4* RNAscope probe was custom designed (Mm-*Abhd4*-O1; #524551) and was used in tandem either with Glast1 (Mm-*Slc1a3*-C2; #430781-C2) or with Tbr2 (Mm-*Eomes*-C2; #429641-C2) to visualize radial glia progenitor cells or intermediate progenitor cells, respectively. To enhance the membrane signal after RNAscope, sections were fixed with 10% PFA for 10 minutes at room temperature. After several, but brief rinsing steps with PBS, sections were incubated with goat antibody to green fluorescent protein (GFP, 1:1000, Abcam) and 5% NDS (Normal Donkey Serum, Sigma) solution at 4°C overnight. Next day, slides were washed with PBS and treated in secondary antibody solution (Alexa Fluor 488-conjugated anti-goat (1:400, Jackson). The multiplex RNAscope signals were imaged by confocal microscopy in randomly selected GFP-expressing cells. The fluorescent dots representing individual mRNA molecules were quantified by manual counting for each marker, and were normalized to the respective median value.

### Western blotting

*Abhd4*-transfected HEK293 cells (as positive controls) and the telencephalon of wild-type and *Abhd4*-knockout embryos (as negative controls) were homogenized in RIPA lysis buffer containing 50 mM Tris-HCl, 150 mM NaCl, 1% Triton X-100, 0.1% SDS, 1mM DTT and 1X protease inhibitor cocktail (Roche). Cytosolic fractions of the samples were denatured in Laemmli sample buffer (Bio-Rad) for 5 minutes at 95°C, and were loaded into a SDS-polyacrylamide gel (12%). Approximately 15 μg of total protein was separated in each lane at 160 V, 400 mA with PowerPac HC High-Current Power Supply and electrophoretically transferred to nitrocellulose membrane (Bio-Rad). To block non-specific binding in immunoblotting, 5% bovine serum albumin (Sigma) was used in Tris-buffered saline containing 0.1% Tween 20 buffer (TBST). The blots were incubated with the rabbit antibody to alpha/beta hydrolase 4 (ABHD4, 1:500, ImmunoGenes) in TBST overnight at 4°C. After several washes with TBST, the membranes were treated with HRP-linked anti-rabbit secondary antibody (1:3000, Cell Signaling) for 2 hours at room temperature and developed by SuperSignal West Dura Extended Duration Substrate Kit (ThermoFisher). Following read-out, the blots were stripped and incubated with the rabbit antibody to catalase (1:3000, Abcam) in TBST overnight at 4°C, and treated as above to confirm comparable protein loading.

The primary antibody against the mouse ABHD4 was raised in transgenic rabbits that have elevated neonatal Fc receptor (FcRn) activity, because they carry one extra copy of the rabbit FcRn α-chain encoding gene (rabbit FCGRT)^58^. The rabbits (3 months old females) were intramuscularly immunized with a keyhole limpet hemocyanin (KLH)-conjugated polypeptide (ABHD4: N′-PNQNKIWTVTVSPEQKDRT-C′) corresponding to amino acid residues 50-69 of the mouse ABHD4 enzyme. Immunization, antiserum selection, affinity purification and antibody validation in knockout mice was performed as described earlier^59^. All the treatments of rabbits in this research followed the guidelines of the Institutional Animal Care and Ethics Committee at ImmunoGenes Ltd that operated in accordance with permissions 22.1/601/000/2009 and XIV-I-001/2086-4/2012 issued by the Food Chain Safety and Animal Health Directorate of the Government Office of Pest County, Hungary. The full Western blot membrane is reported in Supplementary data 2.

### Fluorescence immunostaining and histology

Embryonic brains were immersion fixed with 4% PFA overnight at 4°C. Brains were cryoprotected in 15% sucrose in PBS buffer for 15 minutes and then in 30% sucrose solution overnight at 4°C. After embedding into Tissue-Tek Optimal Cutting Temperature formulation (Sakura), the 20-μm-thick cryosections were collected on Superfrost Ultra Plus glass slides (ThermoFisher) and treated again with 4% PFA for 10 minutes. After several PBS washes, sections were permeabilized with 0.2% Triton X-100 in PBS and were blocked with 1% human serum albumin (Sigma) in PBS for one hour. Primary antibody incubation was carried out overnight at 4°C. The following primary antibodies were used: rabbit antibody to phospho-histone H3 (PHH3, 1:500, Millipore), rabbit antibody to the transcription factor T-box, brain, 1 (TBR1, 1:500, Abcam), rabbit antibody to T-box, brain, 2 (TBR2, 1:500, Abcam), rabbit antibody to laminin subunit alpha 1 (LAMA1, 1:500, Sigma). In case of paired box protein-6 (PAX6)-immunostaining, sections were also treated after fixation with 10 mM citric acid for one hour at 65°C, then washed in 0.1% Triton X-100 in PBS before the blocking step and the primary antibody incubation with rabbit antibody to PAX6 (1:300, Biolegend). In case of BrdU labeling, the sections were treated with 2 M HCl for 1 hour at 37°C and then were blocked with 1% human serum albumin for 1 hour before incubation with the mouse antibody to BrdU (1:1000, Sigma) overnight at 4°C. The following day, the sections were washed extensively in PBS buffer, and then incubated with the appropriate commercial Alexa Fluor 488-conjugated anti-mouse or anti-rabbit (1:400, Jackson), or Alexa Fluor 594-conjugated anti-rabbit (1:400, Jackson) secondary antibodies for 4 hours at room temperature. To visualize F-actin-enriched adherens junctions in the ventricular wall, sections from the embryonic brains were treated as above, but antibody incubation was replaced by the high-affinity F-actin probe Alexa Fluor 568-Phalloidin (1:500, ThermoFisher) treatment for two hours at room temperature. To visualize cell nuclei, DAPI was included in the secondary antibody staining solution according to manufacturer’s protocol (1: 1000 dilution of the 5 mg/ml stock, Calbiochem). Cell death was detected by terminal deoxynucleotidyl transferase dUTP nick end labeling (TUNEL) assay by using Apoptag Red In Situ Apoptosis Detection Kit according to manufacturer’s protocol (Millipore). As the last step after each immunostaining and histological procedures, the slides were washed with PBS, mounted with Vectashield Hardset Antifade Mounting Medium (Vector), covered and sealed with nail polish.

In case of STORM super-resolution imaging, the embryonic brains were fixed in a similar manner, but were cut into thinner 20-µm-thick sections for better signal-to-noise ratio during imaging. Immunostaining was carried out in a free-floating manner. Sections were permeabilized with 0.2% Triton X-100 in PBS and were blocked with 5% normal donkey serum (Sigma) in PBS for one hour. Sections were incubated with a mouse antibody to nestin (1:200, Millipore) overnight at 4 °C, followed by intensive washing in PBS and then incubation with Alexa Fluor 647-conjugated anti-mouse (1:400, Jackson) secondary antibody for 4 hours. Instead of using glass slides, the sections were dried onto coverslips, and stored uncovered at 4°C until STORM imaging.

### In vitro experiments

HEK293 cells were a kind gift from Balázs Gereben (Institute of Experimental Medicine). Cells were maintained under routine conditions in plastic Petri dishes in Dulbecco’s Modified Eagle Medium (4.5 g/L glucose, L-glutamine and sodium pyruvate; Corning) with 10% heat-inactivated fetal bovine serum (Biosera) in a 5% CO_2_ atmosphere at 37 °C. Cells were seeded one day before transfection on poly-D-lysine-coated coverslips in 24-well culture plates. The cells were held in Opti-MEM Media (Gibco) for one hour, and then transfected with 1 μg plasmid DNA in Opti-MEM Media mixed with 2 μl Lipofectamine 2000 Reagent according to manufacturer’s protocol (Invitrogen). After incubation for 20 hours, the transfected cells were washed with PBS and then fixed with 4% PFA for 10 minutes. Subsequently, fixed cells were treated with 0.2% Triton X-100 for 15 minutes at room temperature and blocked with 1% human serum albumin in PBS for 30 minutes. The following primary antibodies were used for immunostaining: rabbit antibody to mitochondrial import receptor subunit TOM20 homolog (TOM20, 1:1000, Santa Cruz), mouse antibody to cytochrome c (CytC, 1:2000, Biolegend), and rabbit anti-cleaved-caspase 3 (CC3, 1:500; Cell Signaling). Primary antibodies were applied in PBS buffer for 1.5 hours at room temperature. After washing steps, the cells were further incubated in CF568-conjugated anti-rabbit (1:1000, Biotium) secondary antibody and Alexa Fluor 647-conjugated anti-mouse (1:400 Jackson) secondary antibody in PBS buffer for one hour at room temperature, and then rinsed extensively with PBS. In case of CC3-immunostaining, the coverslips were mounted with Vectashield Hardset Mounting Medium for confocal microscopy, whereas in case of the dual TOM20- and CytC-immunostaining, the sections were covered in imaging medium for STORM super-resolution microscopy.

### Microscopy

Light micrographs were taken with an Eclipse 80i upright microscope equipped with a DS-Fi1 CCD camera (Nikon). High-resolution fluorescence images were obtained with an A1R confocal laser-scanning system built on a Ti-E inverted microscope and operated by NIS-Elements AR software (Nikon). STORM super-resolution images and the correlated high-power confocal stacks were acquired via a CFI Apo TIRF 100× objective (1.49 NA) on a Ti-E inverted microscope equipped with an N-STORM system, a C2 confocal scan head (Nikon), and an iXon Ultra 897 EMCCD camera (Andor). The HEK293 cells and the embryonic brain sections were covered with a freshly prepared imaging medium containing 0.1 M mercaptoethylamine and components of an oxygen scavenging system consisting of 5% (m/v) glucose, 1 mg/ml glucose oxidase and 2.5 μl/ml catalase in Dulbecco’s PBS (Sigma). The coverslips were sealed with nail polish and transferred into the microscope setup after 10 minutes. Correlated confocal and STORM image acquisition was done as described earlier^59^. In case of HEK293 cells, confocal z-stacks were taken from GFP-expressing cells and TOM20-positive mitochondria were selected to establish the best focal plane for STORM imaging. In case of brain sections, the GFP-positive radial processes of radial glia progenitor cells and the nestin-immunopositive intermediate filaments were located in the cortical plate in confocal z-stacks to determine the optical plane for subsequent STORM imaging. Continuous illumination with 405 nm laser line was used in order to reactivate the fluorophores and produce enough localization events, and 2,500-5,000 cycles were captured using 647-nm excitation. To acquire coordinates of localization points, peak detection was done using the Nikon N-STORM module in the NIS-Elements AR software.

### Image analysis

To analyze correlated confocal and STORM images, pixel-based confocal images and 3D-coordinates of molecular localizations were loaded and aligned in the VividSTORM software^60^. To characterize the 3D-nanoarchitecture of nestin filaments, the number and density of localization points were calculated in randomly selected and identically sized filament segments as region-of-interests. After fitting a convex hull onto the outermost localization points, the volume and surface area of the filaments were also measured. Intensity profiles perpendicular to the nestin filaments were measured to calculate full-width-at-half-maximum values by fitting a Gaussian function in the NIS-Elements software. Gaussian rendering in the N-STORM module was used to visualize higher accuracy localizations by brighter dots. Convex hull and 3D-rendering images were obtained by the Visual Molecular Dynamics software. To investigate mitochondria and the nanoscale distribution of cytochrome c in HEK293 cells, a freehand shape was first drawn around randomly selected GFP-expressing cells as region-of-interests. By using a custom Python script, the localization points representing cytochrome c were counted over the TOM20-positive and TOM20-negative pixels to establish cytochrome c distribution inside and outside of mitochondria, respectively.

Confocal microscopy was used to quantify cellular distribution, morphology and cell death. In distribution analysis, the developing neocortex was divided into five laminar bins and the percentage of GFP-expressing electroporated cells was established in each bin by ImageJ. In morphological analysis, GFP-expressing cells were selected with ImageJ cell counter, rounded cells were counted manually and their percentage was determined. In cell death analysis, TUNEL-positive cell density was measured in the ventricular and subventricular zones with ImageJ cell counter. Cleaved caspase-3-positive and GFP-expressing cells were quantified with NIS-Elements Software.

The density of pyramidal cells, interneurons and astrocytes in the adult neocortex was quantified by Image J/Fiji. Images were auto-thresholded by the following functions: *vGlut1*-Otsu’s; *Gad67*-Li’s; *Glast1*-Triangle and converted to binary image. Pixel numbers with signal were determined in five equal bins and their percentages were counted.

### Prenatal ethanol exposure experiments

Females heterozygous for the *Abhd4* gene were bred with heterozygous males to ensure that littermate wild-type and *Abhd4*-knockout embryos are identically exposed to ethanol *in utero*. In the acute model, pregnant dams received a single intraperitoneal injection of either vehicle or 5g/kg ethanol in saline, and embryos (E14.5) were fixed 12 hours later. In the subchronic model, vehicle or 2.5g/kg ethanol in saline was intraperitoneally injected twice a day between E13.5-E15.5, and embryos were collected at E16. Maternal blood ethanol content was determined enzymatically by using Synchron Systems Ethanol assay kit (Beckman). Blood ethanol standards were created (0.1‰; 0.5‰; 1‰; 1.5‰; 2‰). The dams’ blood alcohol levels were between 0.5‰-1‰ in the subchronic model, whereas it reached 1.5‰-2‰ in the acute model.

### Phylogenetic tree

Protein sequences from different phylogenetic levels were collected from UniProt (https://www.uniprot.org/) and NCBI (https://www.ncbi.nlm.nih.gov/) (accession numbers: Q8TB40, H2Q7Z0, Q8VD66, D3ZAW4, G1T725, Q5EA59, NP_001017287.1, NP_001017613.1, Q8WTS1, Q7JQU9, H2KZ86, A7S6S7). Sequence similarities were determined using protein alignment by Mega7 software^61^. The evolutionary history was inferred by the Maximum Parsimony method. The tree was obtained using the Subtree-Pruning-Regrafting algorithm.

### Statistical analysis

Experimental results were tested for statistical significance using Statistica 13.1 (TIBCO) and Prism 5 (GraphPad). All treatments in each experiment were replicated at least in 3 independent cases. Shapiro-Wilk normality test was used to measure normality of the samples. Unpaired comparisons were analyzed using two-sided Student’s t-tests (normally distributed) and by Mann–Whitney U tests (not normally distributed). Multiple comparisons were tested based on their parametric or non-parametric data by one-way ANOVA with post hoc Tukey’s test or Kruskal-Wallis test followed by post hoc Dunn’s test, respectively. Spearman’s rank correlation coefficient was used to assess relationship between mRNA levels in the single cell-specific RNAscope experiments. All statistical information are reported in Supplementary data 1.

### Data and Code availability

All data, quantitative analysis and scripts are available from the corresponding author.

**Step-by-step Protocols** will be deposited to the open resource Protocol Exchange of Nature Research upon publication.

**Supplementary information** is available for this study.

## Acknowledgements

The authors thank Dr. E. Horváth, B. Pintér, E. Tischler for laboratory support, Drs. G. Balogh, Z. Balogi, A. Dorning, B. Dudok, M. Péter, L. Vígh for help with preliminary experiments and comments. The authors are also grateful to the IEM Medical Genetics Unit for mouse colony management, to Sherry Shu Jung Hu, Kata Nagy for antibody generation, to Dr. B. Gereben for providing HEK293 cells and to Kata Balogh for artwork. The help of Dr. László Barna, the Nikon Microscopy Center at the Institute of Experimental Medicine, Nikon Europe B.V., Nikon Austria GmbH and Auro-Science Consulting is acknowledged for kindly providing microscopy support. This work was supported by the National Research, Development and Innovation Office, Hungary (VKSZ-14-1-2O15-0155 for antibody generation; VEKOP-2.3.3-15-2016-00013 for super-resolution microscopy development and K116915 for ABHD4 research to I.K.). Additional support was provided by the Semmelweis University Grant (EFOP-3.6.3-VEKOP-16-2017-00009 to Z.I.L.), the US National Institutes of Health Grant (DA021696 and DA011322) to K.M. and (DA037660) to B.F.C.

## Author Contributions

I.K., Z.L. and K.M. conceived the project and designed the experiments. Z.I.L. and Z.L. carried out molecular cloning, in situ hybridization, immunostaining, confocal microscopy and data analysis. Z.I.L. carried out western blots, phylogenetic analysis and the in vivo experiments, Z.I.L. and F.M. performed the in vitro experiments. M.Z., V.M. and F.M made STORM imaging and data analysis. G.M.S and B.F.C provided the *Abhd4-*knockout mouse line, I. Kacskovics and K. M. developed and Z.I.L. validated ABHD4 antibodies. Z.L. and I.K. wrote the manuscript with the help of all co-authors.

---

**Supplementary Fig. 1.**
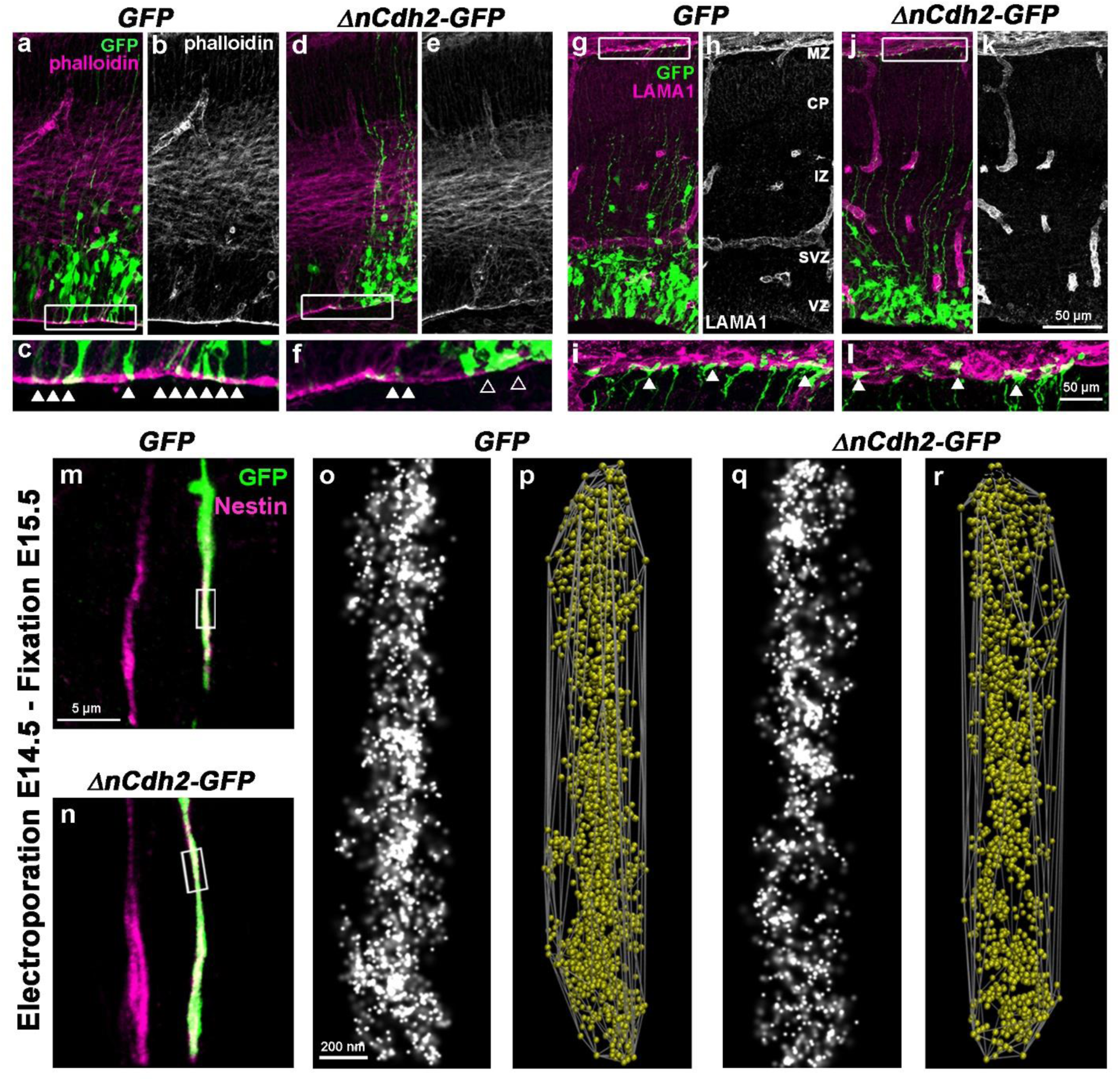
Selective damage of the apical adherens junctions, but not the basal endfeet or the radial processes of RGPCs. **a-f,** Low- (**a,b,d,e**) and high (**c,f**)-power confocal images of the embryonic neocortex stained with the F-actin marker phalloidin. MZ, marginal zone; CP, cortical plate; IZ, intermediate zone; SVZ, subventricular zone; VZ, ventricular zone. Note that *ΔnCdh2-GFP*-electroporation specifically dismantles the adherens junction belt around the affected apical endfeet of RGPCs (**f**), whereas adjacent areas remain intact (white arrowheads), as in control experiments (**c**). **g-l,** Laminin (LAMA1)-immunostaining of the developing cerebral cortex from *GFP*- and *ΔnCdh2-GFP*-electroporated embryos. High-power images (**i,l)** demonstrate that the attachment of the basal endfeet of RGPCs to the pial surface is not affected directly by the disruption of cadherin-based adherens junctions at the ventricular surface. **m-r,** The nanoarchitecture of radial glia processes remains intact. Confocal images of nestin-immunostaining in *GFP-* (**m**) and *ΔnCdh2-GFP-*electroporated (**n**) radial processes. STORM super-resolution images of nestin filaments (**o,q**), and 3D-convex hulls fitted on the nestin localization points to visualize the intermediate filament architecture (**p,r**).

**Supplementary Fig. 2.**
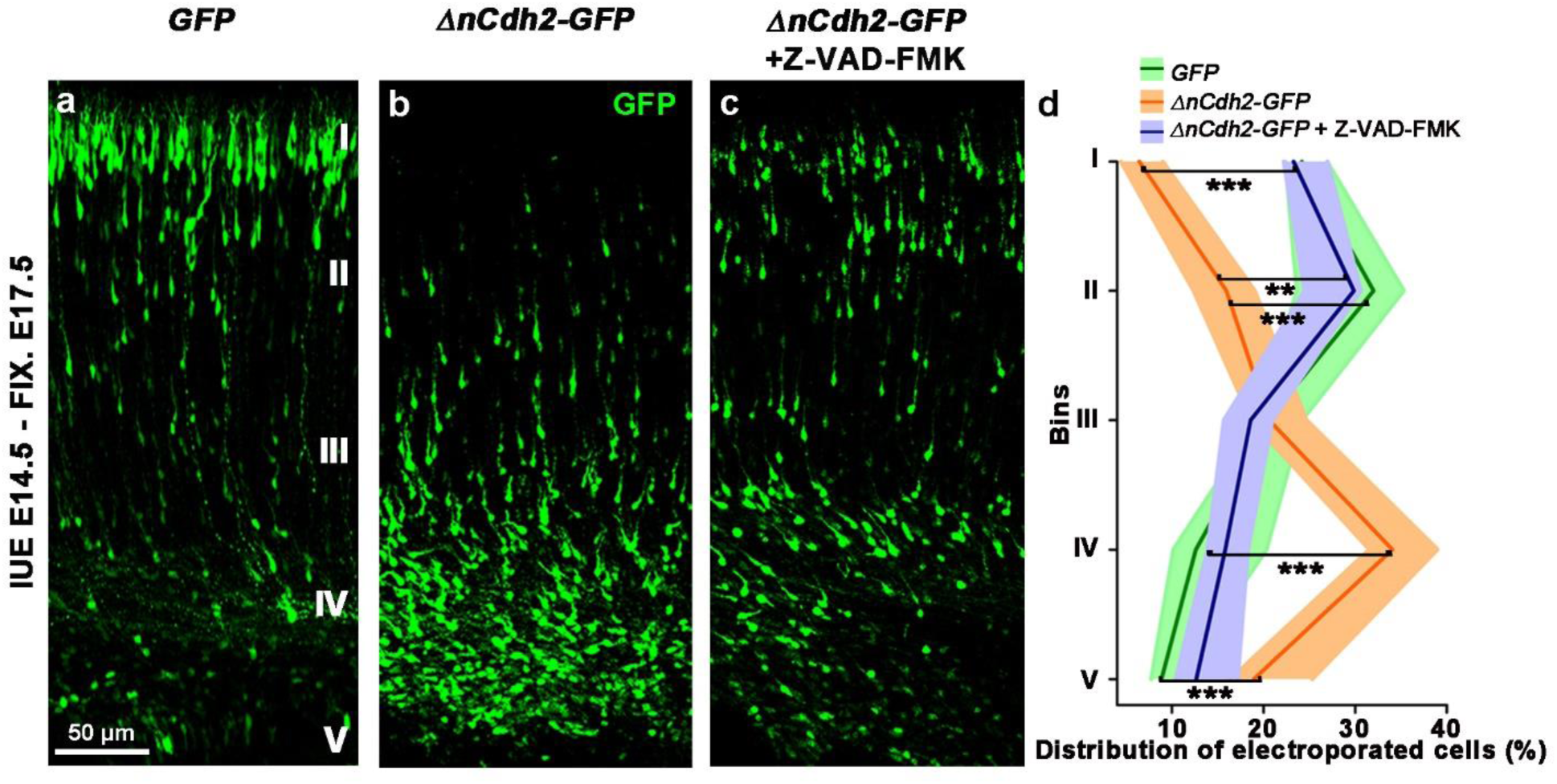
Loss of adherens junctions causes a migration defect. **a,b,** In contrast to migration to the cortical plate under control conditions (**a**), abnormally delaminated cells are concentrated within the subventricular zone (**b**). **c,** Notably, preclusion of cell death by the pan-caspase-inhibitor Z-VAD-FMK treatment restores radial migration. **d,** Quantification of electroporated cells in five equally-sized bins (Roman numerals) (Kruskal-Wallis test with post hoc Dunn’s test, 1^st^,4^th^ bin (*GFP* vs *ΔnCdh2-GFP*, *ΔnCdh2-GFP* vs *ΔnCdh2-GFP +* Z-VAD-FMK): \*\*\**P <* 0.0001; 5^th^ bin (*GFP* vs *ΔnCdh2-GFP*, *ΔnCdh2-GFP* vs *ΔnCdh2-GFP +* Z-VAD-FMK): \*\*\**P =* 0.0001; 2^nd^ bin (*GFP* vs *ΔnCdh2-GFP):* \*\*\**P =* 0.0002; 2^nd^ bin (*ΔnCdh2-GFP* vs *ΔnCdh2-GFP +* Z-VAD-FMK ): \*\**P =* 0.0031; *n* = 10 sections from *n* = 3 animals per *GFP* treatment; *n* = 12 sections from *n* = 3 animals per *ΔnCdh2-GFP* treatment; *n* = 14 sections from *n* = 3 animals per *ΔnCdh2-GFP +* Z-VAD-FMK treatment). Data are shown as median (line) and interquartile range (transparent band in the same color).

**Supplementary Fig. 3.**
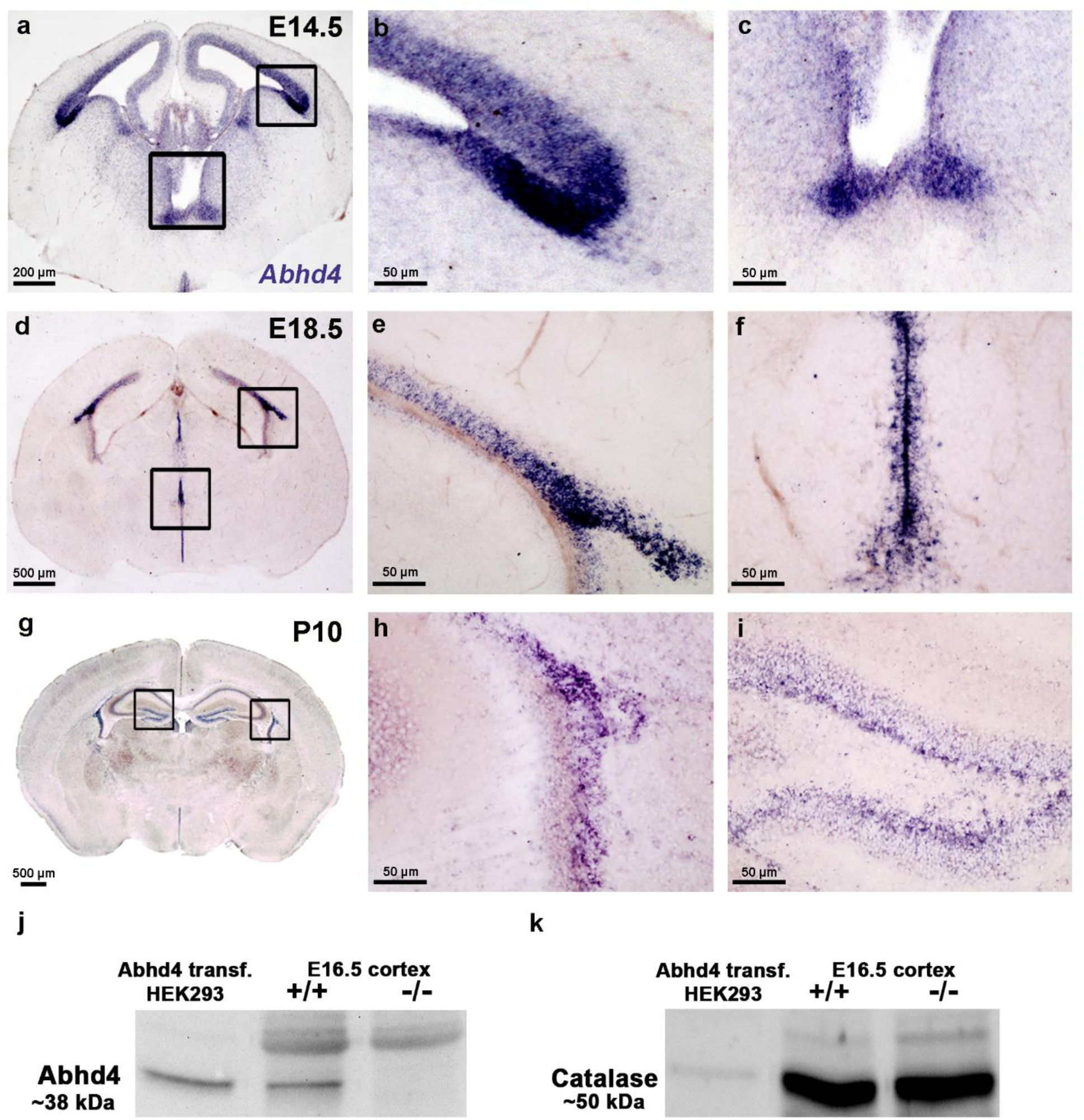
Spatiotemporally uniform *Abhd4* expression pattern in the germinative niches of the developing brain. **a-i,** Coronal sections through the forebrain during prenatal and early postnatal development. Low-power light micrographs taken at embryonic days E14.5 (**a**) and E18.5 (**d**) and postnatal days P10 (**g**) highlight the consistently high expression of *Abhd4* mRNA in the proliferative zones throughout development. High-power light micrographs obtained from the lateral (**b,e**) and the third ventricle (**c,f**) and from the subventricular (**h**) and subgranular layers (**i**) demonstrate that Abhd4 mRNA levels are under the detection threshold outside of the germinative niches. **j,** Western blots verify ABHD4 protein in control *Abhd4-GFP*-transfected HEK293 cells and in the developing neocortex of wild-type (+/+), but not *Abhd4-*knockout mice at E16.5. **k**, Immunoblotting for catalase is used as loading control.

**Supplementary Fig. 4.**
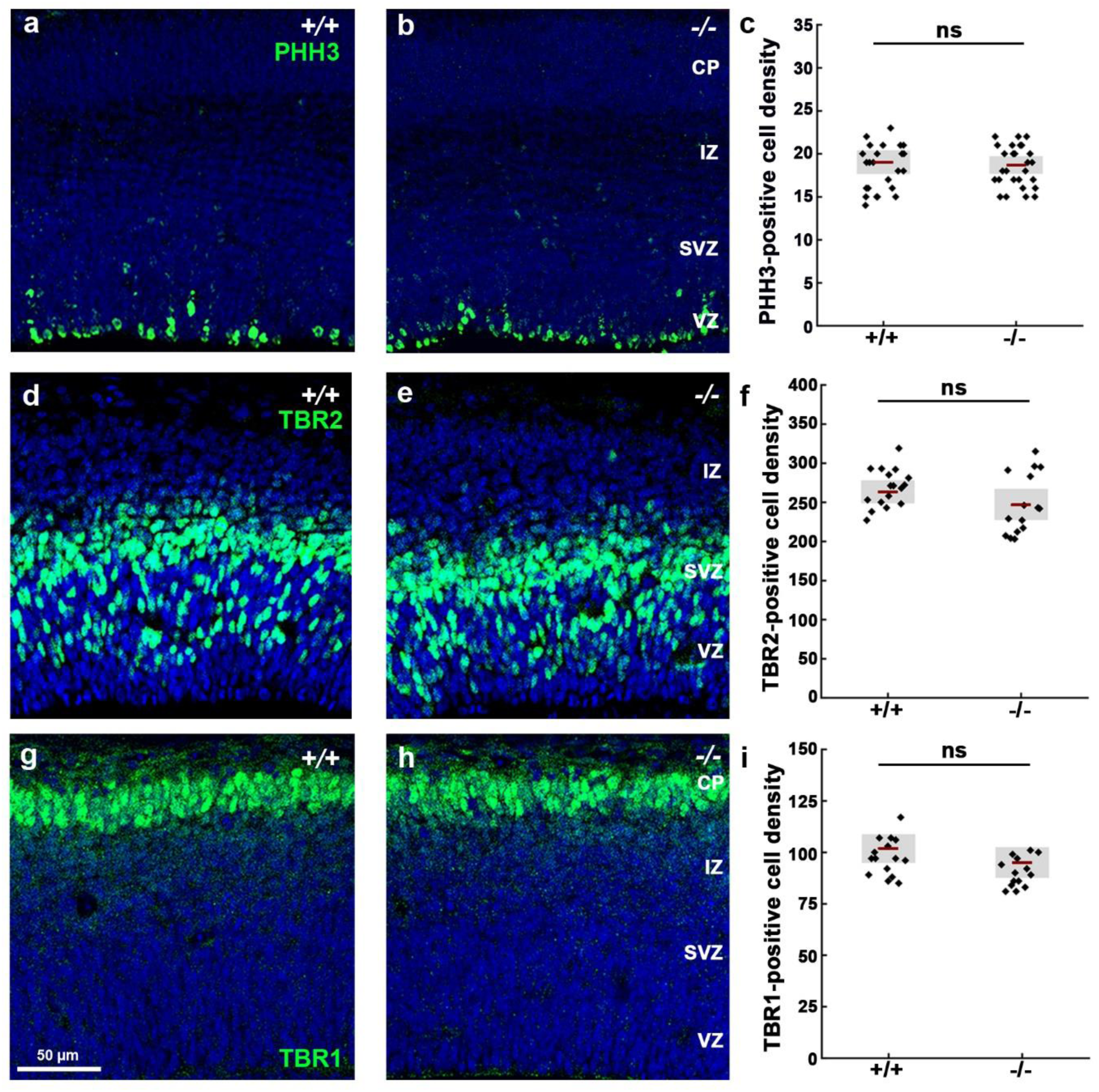
ABHD4 is not required for proliferation, differentiation and lamination in the developing neocortex. **a,b,** Phospho-histone H3 (PHH3)-immunostaining visualizes cell proliferation during the M phase of the cell cycle at the ventricular border. **c,** Quantification of PHH3-positive cell density (two-sided Student’s unpaired t-test, *P =* 0.688; *n =* 26 sections from *n =* 3 animals per wild-type (+/+) mice, *n =* 30 sections from *n =* 3 animals per *Abhd4-*knockout (-/-) mice). **d,e,** High density of T-box brain protein 2 (TBR2)-positive intermediate progenitor cells in the subventricular zone. **f,** Quantification of TBR2-positive cell density (two-sided Student’s unpaired t-test, *P =* 0.194; *n =* 18 sections from *n =* 3 animals per wild-type (+/+) mice, *n =* 15 sections from *n =* 3 animals per *Abhd4-*knockout (-/-) mice). **g,h**, Distribution of TBR1-positive postmitotic neurons in the cortical plate. **i,** Quantification of TBR1-positive cell density (two-sided Mann-Whitney U test, *P =* 0.074; *n =* 17 sections from *n =* 3 animals per wild-type (+/+) mice, *n =* 16 sections from *n =* 3 animals per *Abhd4-*knockout (-/-) mice). MZ: marginal zone, CP: cortical plate, IZ: intermediate zone, SVZ: subventricular zone, VZ: ventricular zone. Graphs show raw data and mean ± 2 x standard error (**c,f**), or raw data and median ± interquartile range (**i**).

**Supplementary Fig. 5.**
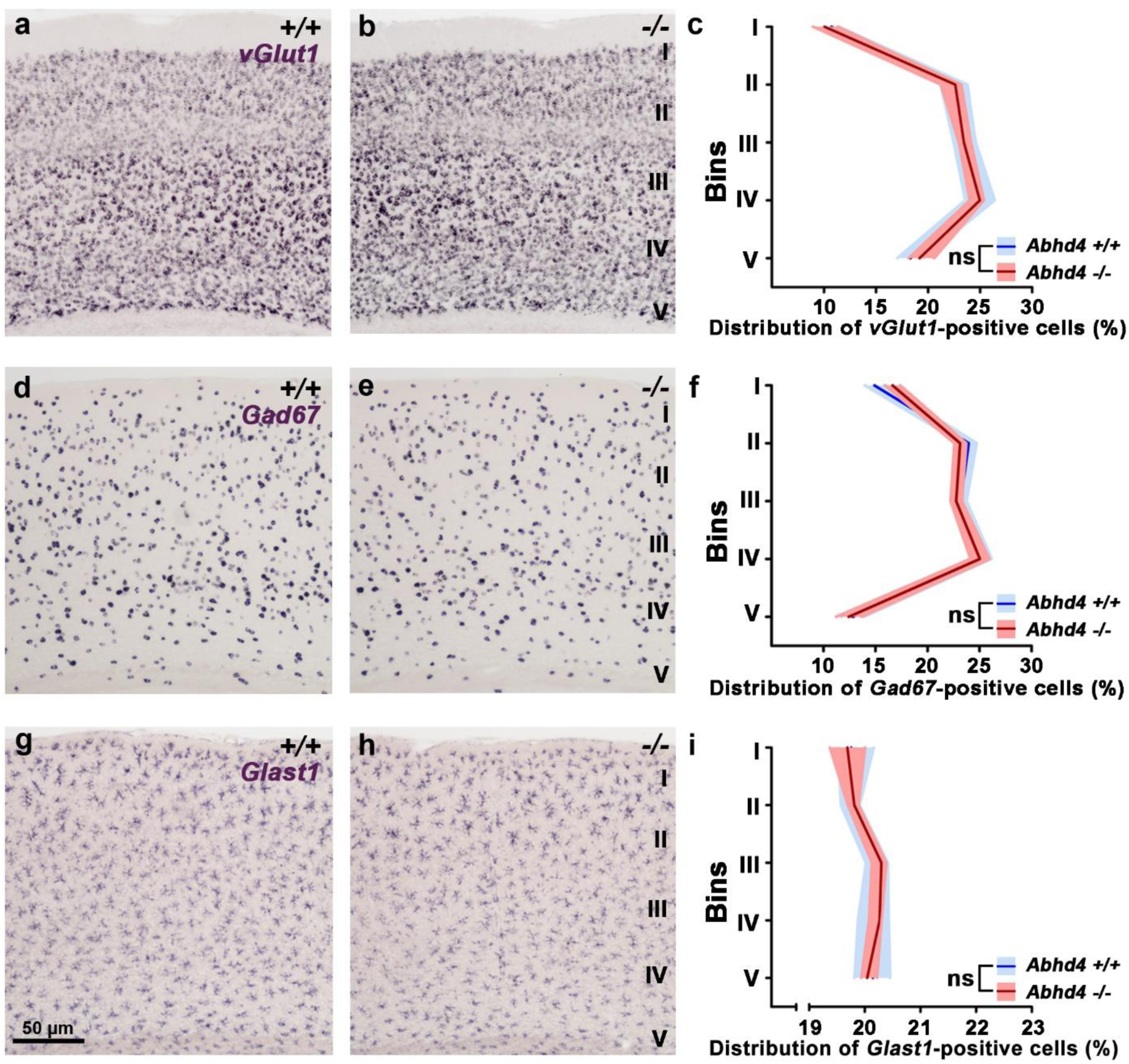
ABHD4 is not necessary for the differentiation and laminar distribution of adult cortical cell types. **a,b,** Distribution of vGlut1-positive excitatory cells in coronal sections of the adult neocortex of wild-type (+/+) and *Abhd4-*knockout (-/-) mice. **c,** Laminar density of *in situ* hybridization signal for *Slc17a7* (encoding the VGLUT1 protein). **d,e,** Distribution of *Gad67*-positive inhibitory cells in coronal sections of the adult neocortex of wild-type (+/+) and *Abhd4-*knockout (-/-) mice. **f,** Laminar density of *in situ* hybridization signal for *Gad1* (encoding the GAD67 protein). **g,h,** Distribution of *Glast1*-positive astrocytes in coronal sections of the adult neocortex of wild-type (+/+) and *Abhd4-*knockout (-/-) mice. **i,** Laminar density of *in situ* hybridization signal for *Slc1a3* (encoding the GLAST1 protein). Density measurements were performed in five equally-sized bins (Roman numerals) (two-sided Mann-Whitney U test, *P >* 0.05 in all bins of all experiments, see accompanying Supplementary data 1. for exact values; *n =* 22 sections from *n =* 3 animals per wild-type (+/+) mice, *n =* 26 sections from *n =* 3 animals per *Abhd4-*knockout (-/-) mice in case of vGlut1; *n =* 23 sections from *n =* 3 animals per wild-type (+/+) mice, *n =* 24 sections from *n =* 3 animals per *Abhd4-*knockout (-/-) mice in case of Gad67 and Glast1). Data are shown as median (line) and interquartile range (transparent band in the same color).

**Supplementary Fig. 6.**
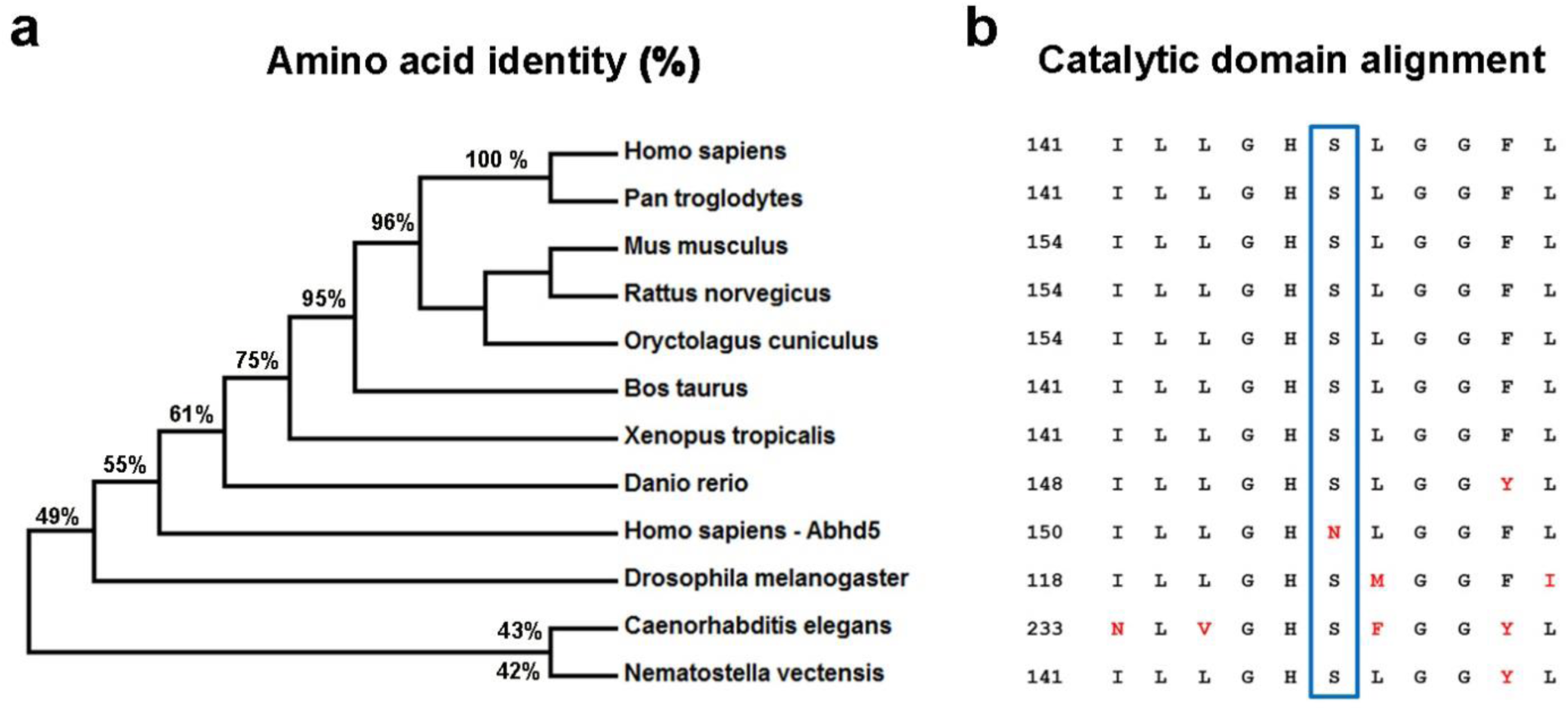
ABHD4 is an evolutionarily conserved serine hydrolase. **a,** The phylogenetic tree of ABHD4 protein sequences. Percentage values indicate the amino acid identity between the human ABHD4 protein sequence and the respective orthologues of other species. Human ABHD5, the closest relative of ABHD4 is also included for comparison. **b,** Note that ABHD4 orthologues exhibit substantial homology that is especially high around the catalytic serine residue (blue box) and the consensus “nucleophile elbow” sequence “GXSXG”. In contrast, ABHD5, the paralogous enzyme that diverged from ABHD4 ∼500 million years ago, lost its catalytic serine during evolution.

**Supplementary Fig. 7.**
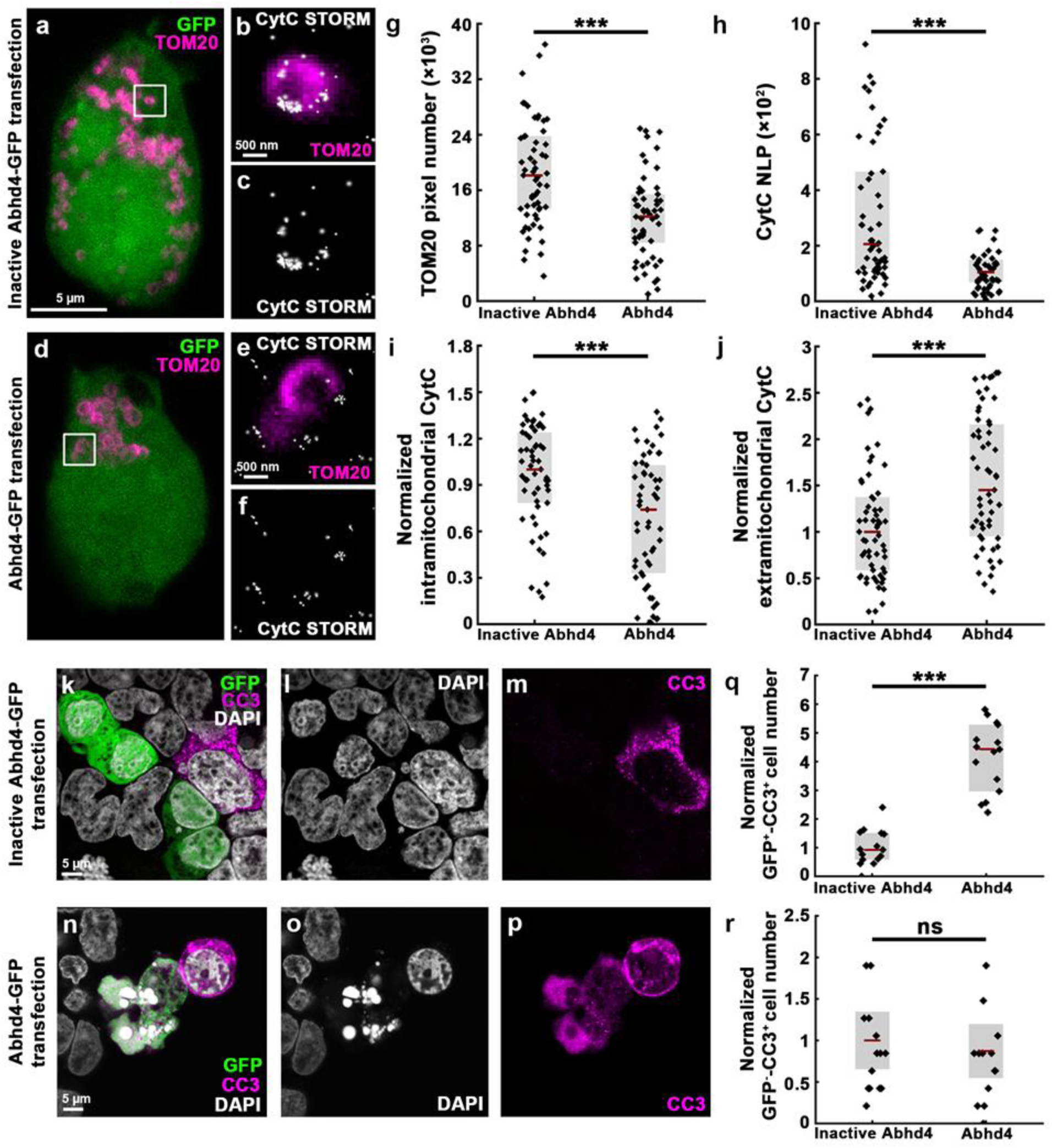
ABHD4 drives apoptosis cell-autonomously. **a-f,** Correlated confocal and STORM super-resolution images of mitochondrial import receptor subunit TOM20 homolog (TOM20)- and cytochrome c (CytC)-immunostaining, respectively, of HEK293 cells transfected either with wild-type (**d**) or with an enzymatically inactive form of *Abhd4-GFP* (**a**). Note that STORM localization points representing individual CytC proteins are concentrated in the TOM20-positive mitochondrion in the transfected HEK293 cell under control condition (**b,c**), but are released into the extramitochondrial environment in the *Abhd4-GFP* electroporated cell (**e,f**). **g,** Quantification of TOM20-immunostaining in transfected HEK293 cells. **h,** The number of localization points (NLP) representing CytC in the same cells. **i**, Normalized number of CytC-localization points within TOM20-positive mitochondria. **j**, Normalized number of CytC-localization points outside of TOM20-positive mitochondria. Two-sided Mann-Whitney U test in **g-j**, \*\*\**P <* 0.0001 in **g,h**; \*\*\**P =* 0.0003 in **i,j**; *n =* 61 cells from *n =* 4 cultures per Inactive *Abhd4-GFP* treatment, *n =* 57 cells from *n =* 4 cultures per *Abhd4-GFP* treatment. **k-p,** Confocal images show cleaved caspase-3 (CC3)-immunostaining in HEK293 cells after electroporation with *Abhd4-GFP* (**k-m**), but not with Inactive *Abhd4-GFP* (**n-p**). Besides the molecular evidence, the characteristic apoptotic signs of cell shrinkage and nuclear fragmentation induced by ABHD4 are also notable (**n-p**). **q,** Quantification of the ratio of GFP-expressing HEK293 cells that are immunostained for CC3 (two-sided Mann-Whitney U test, \*\*\**P <* 0.0001; *n =* 15 cells from *n =* 3 cultures per both treatments). **r,** Statistical analysis of GPF-negative, but cleaved caspase-3-positive normalized cell number (two-sided Student’s unpaired t-test, *P =* 0.5972). Graphs show raw data and median ± interquartile range, except (**r**) which shows mean ± 2 x standard error.

**Supplementary Fig. 8.**
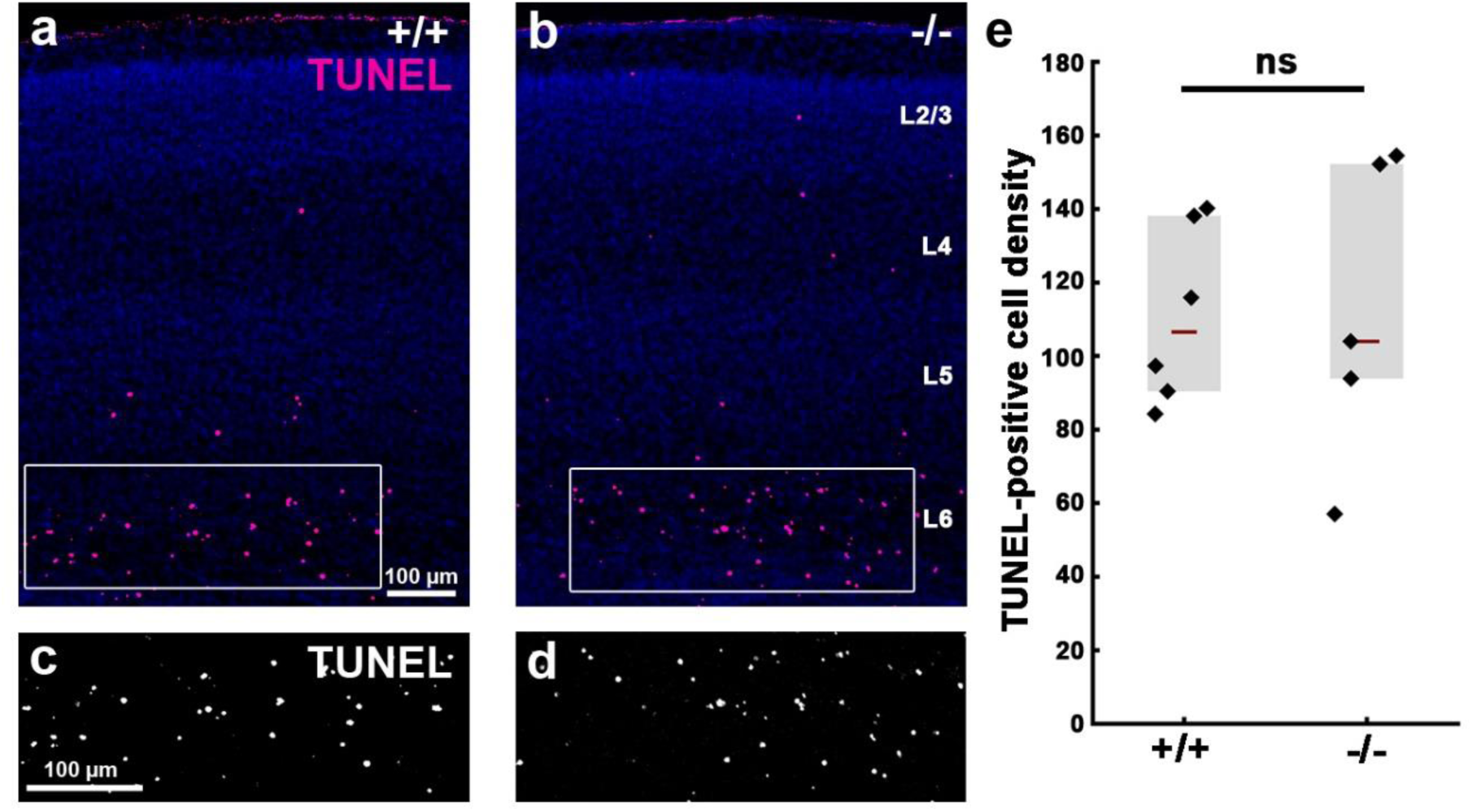
ABHD4 is not required for developmentally controlled programmed cell death in the developing neocortex. **a-d,** Comparable levels of programmed cell death in the early postnatal day 3 (P3) cerebral cortex of wild-type (+/+) and *Abhd4-*knockout (-/-) mice. **e,** Quantification of the density of TUNEL-positive dead cells (two-sided Mann-Whitney U test, *P =* 0.792; from *n =* 6 animals per wild-type (+/+) mice, *n =* 5 animals per *Abhd4-*knockout (-/-) mice). L2/3-6 mark the cortical layers. Data are shown as median ± interquartile range.

**Supplementary Fig. 9.**
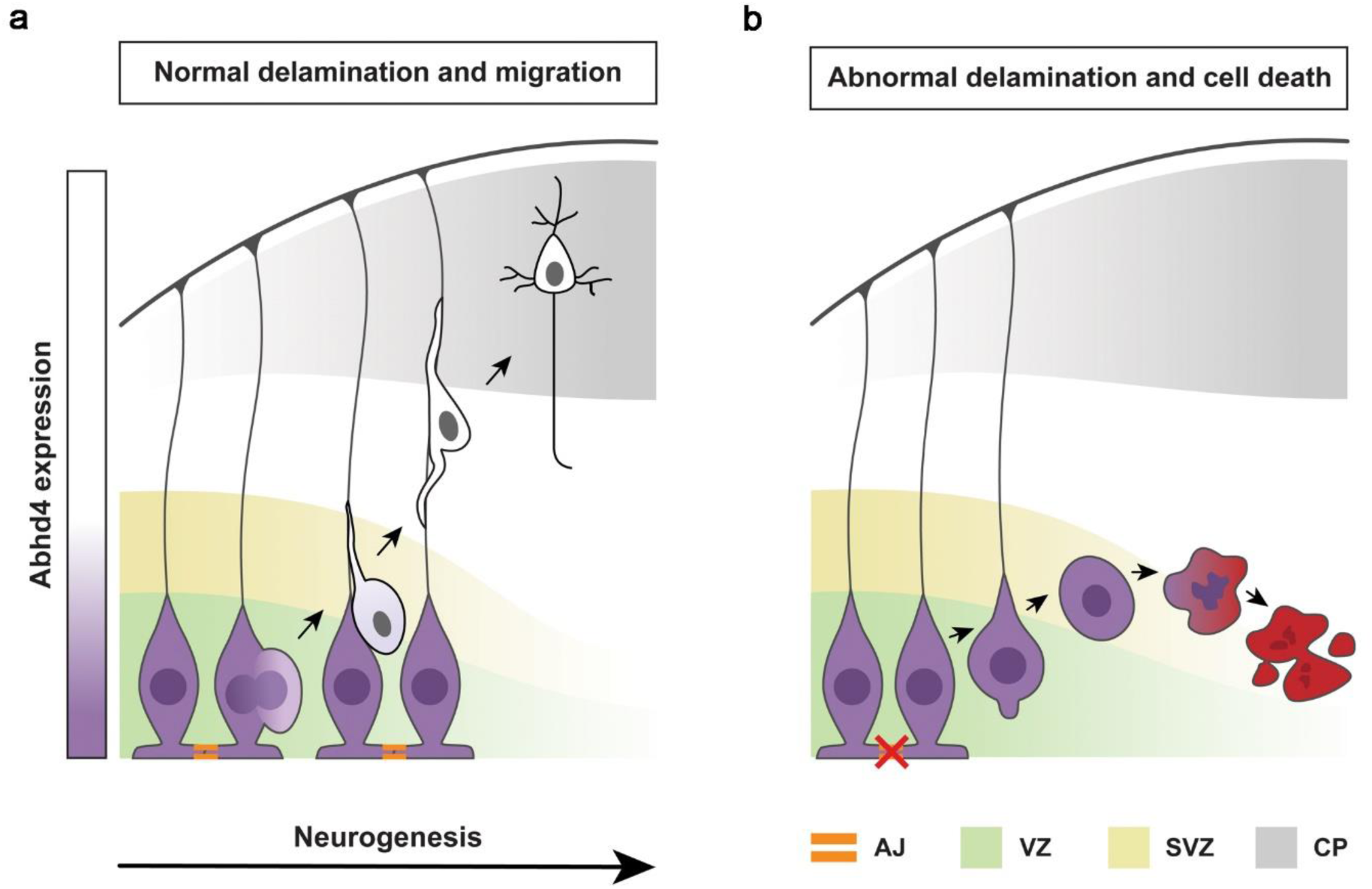
ABHD4 is a molecular mediator of developmental anoikis that distinguishes premature and mature delaminating cells. **a,b**, Schematic iluustrates the distinct ABHD4-dependent fate of normally and abnormally delaminated cells in the developing cerebral cortex. After cell division, healthy daughter cells downregulate ABHD4, delaminate from the ventricular wall and migrate into the cortical plate along the radial processes of RGPCs. Random developmental errors and pathological insults such as fetal alcohol exposure often destroy the adherens junctions and trigger abnormal delamination. As a central mediator of developmental anoikis, ABHD4 is necessary and sufficient to induce cell death that prevents the accumulation of misplaced progenitor cells at ectopic locations. CP, cortical plate; SVZ, subventricular zone; VZ, ventricular zone.

